# Directional heritability and the geometry of multivariate constraint

**DOI:** 10.64898/2026.01.26.701879

**Authors:** Nicholas O’Brien, Pamela Burrage, Kevin Burrage, Daniel Ortiz-Barrientos

**Affiliations:** School of the Environment, The University of Queensland, St Lucia, QLD 4072, Australia; School of Mathematical Sciences, Queensland University of Technology, Brisbane, QLD 4001, Australia; ARC Centre for Plant Success in Nature and Agriculture, The University of Queensland, St Lucia, QLD 4072, Australia

**Keywords:** evolvability, genetic variance-covariance matrix, directional heritability, evolutionary constraint, quantitative genetics

## Abstract

Evolutionary responses to selection depend on how additive genetic variance is distributed across trait combinations. We focus on directional heritability—the fraction of phenotypic variance that is additive genetic along a given selection gradient—and treat its distribution across directions as a central object for describing multivariate constraint. Using a geometric transformation that rescales trait space so that phenotypic variance is isotropic, we show that directional heritability becomes a quadratic form in a whitened genetic matrix **G**^∗^ = **P**^−1/2^**GP**^−1/2^. Under uniformly distributed selection directions, the squared coefficient of variation of directional heritability satisfies CV^2^(*h*^2^) = (2/(*p* + 2)) *V*_rel_(**G**^∗^), where *V*_rel_(**G**^∗^) is the relative eigenvalue variance of the whitened matrix and *p* is the number of traits. Simulations show that alignment between the eigenvector systems of **G** and **P** has a larger effect on the spread of directional heritability than correlation strength. Analyses of 55 empirical **G**–**P** matrix pairs from 11 studies reveal wide variation across biological systems in how often selection encounters low-heritability directions: over two thirds of the populations examined had more than 25% of phenotypic directions with *h*^2^ *<* 0.25. The eigenvalue spectrum of **G**^∗^ provides a sufficient summary for characterising how matrix geometry shapes evolutionary constraint on the phenotypic scale.

## Introduction

Selection in nature rarely pushes in a fixed direction. Climatic oscillations, novel competitors, and shifts in resources continually redirect selective pressures across trait space. A population that evolves readily under one selective regime may be constrained when selection pivots – not because genetic variance is absent, but because existing variance may be poorly aligned with what selection now demands. This raises the question: when selection targets an arbitrary direction in phenotype space, how likely is a population to respond efficiently?

The short-term response to selection on multivariate phenotypes is given by Lande’s equation 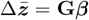 (Lande 1979), where **G** is the additive genetic variance-covariance matrix and ***β*** is the selection gradient. This predicts the direction and magnitude of evolutionary change for a specified ***β***. When **G** contains strong genetic covariances, the response to selection can vary dramatically with ***β***. Specifically, the alignment of ***β*** with *G*_*max*_, the major axis of variation in **G**, determines the magnitude of the response (Schluter 1996). When future selection is uncertain, a population’s evolutionary potential depends not on its capacity to respond to any particular gradient, but on how genetic variance is apportioned across the space of directions that selection might take.

Hansen and Houle (2008) addressed this by introducing evolvability, 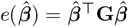, the additive genetic variance in direction 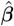. They showed that the average evolvability across random directions is given by *ē* = tr(**G**)*/p*, where *p* is the number of traits in **G**. While *ē* provides a useful scalar summary, as Hansen and Houle (2008) noted, this average can obscure substantial heterogeneity: two populations may share the same *ē* yet differ in how evenly genetic variance is distributed across directions. One architecture may concentrate variance along a few axes with near-zero variance elsewhere; another may spread variance more uniformly. These cases have identical average evolvability but the specific evolutionary outcomes for a given direction could differ greatly.

To characterise this variation on a phenotypic scale, we turn to directional heritability (Lin and Allaire 1977):

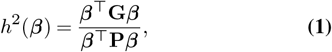

the fraction of phenotypic variance that is additive genetic in direction ***β***. This quantity captures what selection actually “sees”: phenotypic variation, of which only the genetic component transmits to offspring. A direction with substantial genetic variance but even larger environmental variance presents a weak signal-to-noise ratio; selection encounters phenotypic differences yet achieves relatively limited genetic gain (Alatalo et al. 1990). Directional heritability measures this efficiency directly.

Our aim is to describe how *h*^2^(***β***) varies across gradient space and to connect that variation to the geometry of **G** and **P**. We (i) derive an exact relationship linking the coefficient of variation of directional heritability to the eigenvalue distribution of a whitened matrix, (ii) validate this relationship through simulation, and (iii) apply the framework to empirical datasets spanning diverse taxa. Together, these results show that the spread of directional heritability across selection directions is controlled by the eigenvalue dispersion of the whitened matrix **G**^∗^, that alignment between **G** and **P** is a major driver of heterogeneity in *h*^2^(***β***), and that empirical **G**–**P** pairs differ not only in average heritability but also in how often selection encounters low-heritability directions.

## Theory

### Genetic axes, generalised eigenvalues, and null models

We summarise multivariate genetic constraint by solving a generalised eigenvalue problem of the form

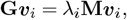

where **G** is the additive genetic covariance matrix, **M** is a scaling matrix (typically the phenotypic covariance matrix **P** or the identity **I**), ***v***_*i*_ is a linear combination of traits (an eigenvector), and *λ*_*i*_ is the associated generalised eigenvalue. Biologically, each ***v***_*i*_ defines a direction in trait space along which the population can evolve, and *λ*_*i*_ measures the heritable variance available along that direction. This becomes evolvability when **M** = **I**, or directional heritability when **M** = **P** (e.g. Houle 1992).

To assess whether observed patterns of constraint are biologically meaningful, we compare eigenvalues and the alignment between eigenvectors and selection directions to null distributions for angles and quadratic forms in high-dimensional spaces. Under these null models, the squared cosine of the angle between a random direction and any fixed axis follows a beta distribution, and evolvabilities along random directions follow distributions that can be written as ratios of quadratic forms (Watanabe 2024). In high-dimensional trait spaces, random directions are nearly orthogonal to any fixed axis, and evolvabilities are relatively narrowly distributed around their mean. The strong concentration of heritability along particular axes, and the large spread of directional heritability across selection directions, therefore cannot be explained by geometry alone. These departures from null expectations (developed in detail in Appendices 1–3) point to biologically meaningful constraints on evolution rather than artefacts of high-dimensional random structure.

### Whitening and the representation of directional heritability

The ratio *h*^2^(***β***) = ***β***^⊤^**G*β***/***β***^⊤^**P*β*** involves two quadratic forms. To simplify analysis, we transform to coordinates where phenotypic variance is isotropic.

Define ***z*** = **P**^1/2^***β***/ ∥**P**^1/2^***β***∥, a unit vector in Euclidean trait space, and the **P**-whitened genetic matrix

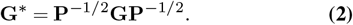

Note that **G**^∗^ is referred to as **G**_**P**_ in Hansen and Houle (2008). A short calculation (Appendix 1) shows that

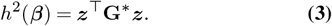

In whitened coordinates, directional heritability is simply a quadratic form evaluated on a unit vector, ***z***. The eigenvalues of **G**^∗^ are the heritabilities along orthogonal directions in whitened space; its eigenvectors define these directions. Figure 1 illustrates this transformation geometrically.

**Figure 1.**
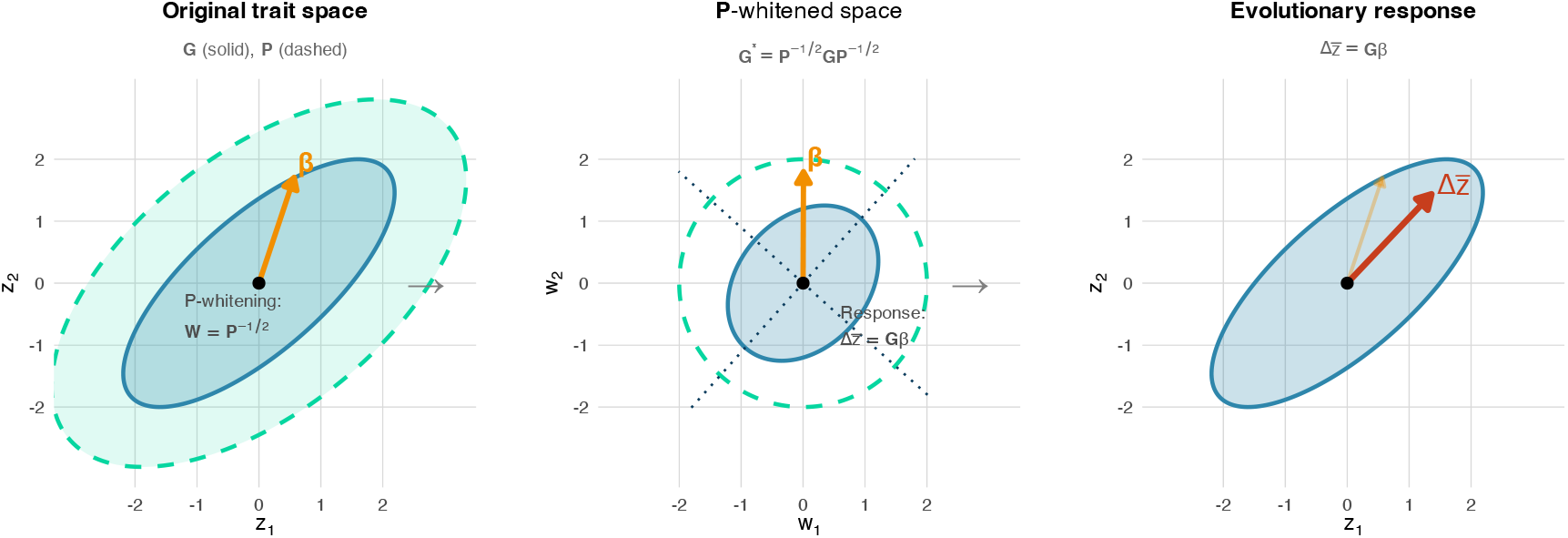
Geometric transformation through P-whitening and evolutionary response. The P-whitening transformation maps the original trait space (left) to a coordinate system where phenotypic variance is isotropic (centre), revealing how the response to selection depends on the interaction between **G** and **P**. *Left panel*: Original trait space showing the genetic variance-covariance matrix **G** (blue solid ellipse) and phenotypic matrix **P** (teal dashed ellipse) with selection gradient *β* (orange arrow). *Centre panel*: After P-whitening (**W** = **P**^−1/2^), the phenotypic variance becomes spherical (dashed circle) while **G** transforms to **G**^∗^ = **P**^−1/2^**GP**^−1/2^ (blue ellipse). The eigenvectors of **G**^∗^ (dotted lines) indicate directions of maximum and minimum heritability. *Right panel*: The evolutionary response 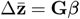 (red arrow) deviates from the selection gradient due to genetic covariances, with the original selection direction shown for reference (faint orange).

#### Box 1: Why We “Whiten” Matrices

If the matrix algebra feels abstract, visualize the population as a cloud of data points. A variance-covariance matrix describes the shape of this cloud: the phenotypic matrix **P** defines an ellipse showing how phenotypic variation is spread, while **G** describes the spread of breeding values.

Directional heritability *h*^2^(***β***) compares the “length” of the genetic ellipse to the “length” of the phenotypic ellipse in direction ***β***: a signal-to-noise ratio. The analytical challenge is that both the numerator and the denominator change as you rotate through ***β*** directions, making the ratio difficult to characterize.

“Whitening” solves this by transforming coordinates. Rescaling trait space by **P**^−1/2^ “un-stretches” the phenotypic ellipse into a perfect circle (or sphere in higher dimensions), similar to how we standardize variables by dividing by their standard deviation, but in this case, in multiple dimensions simultaneously. In whitened space, a unit step in any direction corresponds to exactly one unit of phenotypic variance. The denominator becomes constant and drops out, leaving a single quadratic form: *h*^2^ = **z**^⊤^**G**^∗^**z**.

In the whitened matrix **G**^∗^ = **P**^−1/2^**GP**^−1/2^, eigenvalues are heritabilities along orthogonal directions, and their dispersion determines how much *h*^2^ varies across selection gradients.

Readers unfamiliar with whitening transformations are encouraged to also read Appendix S1 before proceeding.

### Distribution under random selection

If selection directions are uniformly distributed on the **P**-unit sphere, then ***z*** is uniformly distributed on the Euclidean unit sphere *S*^*p*−1^. Writing the eigendecomposition **G**^∗^ = **U**Λ^∗^**U**^⊤^ and ***w*** = **U**^⊤^***z***, we obtain

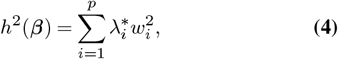

a weighted average of eigenvalues with weights 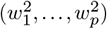 that follow a symmetric Dirichlet distribution with all parameters equal to 1/2. Using the standard moments of this distribution, 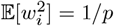 and 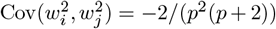 for *I* ≠ *j*, we obtain (Appendix 2):

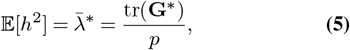

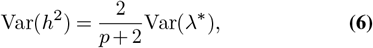

where Var(*λ*^∗^) is the variance of the eigenvalues of **G**^∗^.

Defining the relative eigenvalue variance as 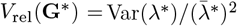, we arrive at our main result:

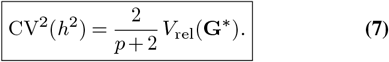

This relationship is exact. It shows that the spread of directional heritabilities across random selection directions depends only on the number of traits (*p*) and the eigenvalue dispersion of the whitened matrix. The factor 2/(*p* + 2) arises from the Dirichlet geometry of random directions and implies that higher dimensionality tends to concentrate directional heritability around its mean. Biological implications of this relationship are identified in Table 1.

**Table 1.** Biological implications of 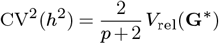.

| Lever | Mathematical effect | Biological implication | Main caveat |
| --- | --- | --- | --- |
| Increase in effective dimensionality $p$ (more independent directions with non-negligible eigenvalues of $\mathbf{G}^*$) | For fixed $V_{\text{rel}}(\mathbf{G}^*)$, $CV(h^2)$ shrinks roughly as $1/\sqrt{p}$ via the factor $\sqrt{2/(p+2)}$. | Heritability tends to become more uniform across directions; random selection is less likely to fall into low-heritability directions. In unpredictable environments, architectures that maintain many moderately heritable directions may be favoured because they reduce constraint events. | Adding traits does not help if most new eigenvalues are near zero. Effective dimensionality, not raw trait count, matters. |
| Increase in $V_{\text{rel}}(\mathbf{G}^*)$ (more unequal whitened eigenvalues) | For fixed $p$, $CV(h^2)$ increases linearly with $\sqrt{V_{\text{rel}}(\mathbf{G}^*)}$. | Heritable signal becomes more uneven across directions: some trait combinations are far more heritable than others. This suits stable or recurrent selection along a few axes, but raises the risk that novel selection falls in poorly heritable directions. | High $V_{\text{rel}}(\mathbf{G}^*)$ can be beneficial when selection is predictable but may be costly when it is diffuse or fluctuating. |
| Strong G–P alignment (eigenvectors of $\mathbf{G}$ and $\mathbf{P}$ nearly coincide) | Alignment tends to reduce $V_{\text{rel}}(\mathbf{G}^*)$ for a given pair of raw spectra, because large genetic and phenotypic eigenvalues align. This lowers $CV(h^2)$. | When genetic variance is concentrated where phenotypic variance is also high, heritability tends to be more stable across directions. This can explain cases where extreme eigenvalue ratios in $\mathbf{G}$ coexist with modest variation in directional heritability. | Maintaining strong alignment may be difficult as trait number and pleiotropy increase, especially under changing environments. |
| Strong G–P misalignment | Misalignment can inflate $V_{\text{rel}}(\mathbf{G}^*)$ even when $V_{\text{rel}}(\mathbf{G})$ and $V_{\text{rel}}(\mathbf{P})$ are moderate, raising $CV(h^2)$. | Can create directions with substantial phenotypic variance but relatively little additive genetic variance. In fluctuating environments this increases the chance that selection sometimes acts in poorly heritable directions. | Misalignment arises from environmental noise, developmental structure, or historical selection; selection can reduce but not necessarily eliminate it. |
| Modular genetic architecture (blocks of traits with strong within-module, weak between-module covariances) | Within modules, effective $p$ is modest and $V_{\text{rel}}(\mathbf{G}^*)$ can be kept low; across modules, the organism behaves like a higher-dimensional system with several moderately isotropic blocks. | Modularity can combine the benefits of higher dimensionality (many directions with usable heritability) with relatively low $V_{\text{rel}}$ within functional sets of traits. This may maintain heritable signal across modules while limiting cross-module constraints. | The evolution of modularity depends on recombination, pleiotropy and the structure of selection; trade-offs with integration and performance in specific environments remain system-specific. |
| Increasing trait number without changing architecture (more traits but same pleiotropic pattern) | Raw $p$ increases, but if most new directions have tiny eigenvalues in $\mathbf{G}^*$, effective dimensionality changes little; $V_{\text{rel}}(\mathbf{G}^*)$ may even rise. | Measuring more traits does not automatically stabilise directional heritability. If new traits are weakly heritable or tightly constrained, the organism still experiences high variation in $h^2(\beta)$ across selection directions. | Highlights that Eq. 7 concerns the geometry of whitened eigenvalues, not trait labels; empirical work should distinguish between measured and effective dimensionality. |

### Constraint probability

For a threshold *c* ∈ [0, 1], we define the constraint probability (or “constraint risk”)

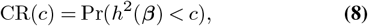

the probability that a randomly oriented selection gradient encounters directional heritability below *c*. This quantity translates eigenvalue geometry into a statement about evolutionary risk: how often would selection, if applied in random directions, encounter low-heritability regions of phenotype space? In practice, we use thresholds such as *c* = 0.25 as benchmarks for low heritability, but the framework is agnostic about the particular choice of *c*.

## Simulation Study

The theory links the spread of directional heritability under a simple baseline for selection directions to the eigenvalue dispersion of the whitened matrix **G**^∗^. The simulation study has two aims. First, it provides a numerical check of Eq. 7 under the same sampling geometry assumed in the derivation. Second, it illustrates how familiar features of genetic architecture—eigenvalue dispersion, correlation structure, and **G**–**P** misalignment—map onto variation in *h*^2^(***β***) on the phenotypic scale.

We worked in *p* = 5 trait dimensions and generated synthetic positive-definite pairs (**G, P**) spanning a range of eigen-value spectra and relative orientations. For each matrix pair we sampled many selection directions under the uniform-on-**P**-sphere baseline, computed *h*^2^(***β***) for each direction, and summarised the resulting distribution using CV(*h*^2^) and the low-tail mass (constraint probability) at an illustrative threshold. To match the theoretical setup, we implemented sampling by drawing ***z*** uniformly from the Euclidean unit sphere and evaluating *h*^2^ = ***z***^⊤^**G**^∗^***z***, where **G**^∗^ = **P**^−1/2^**GP**^−1/2^.

Across the sweep, Monte Carlo estimates of CV^2^(*h*^2^) tracked the theoretical prediction from Eq. 7 (Figure 2). This agreement is expected given the derivation, but it is still useful as a check that the whitening, eigendecompositions, and sampling geometry were implemented consistently.

**Figure 2.**
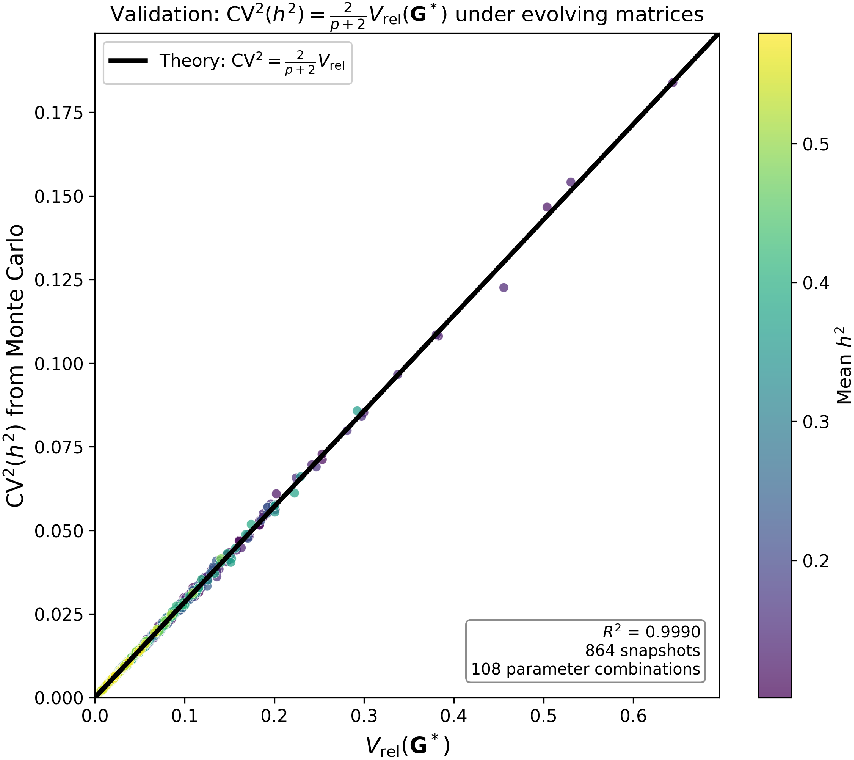
Numerical check of Eq. 7. Each point is a matrix snapshot from the simulation sweep. Monte Carlo estimates of CV^2^(*h*^2^) fall on the theoretical prediction 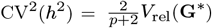.

The sweep also helps organise intuition about what drives heterogeneity in *h*^2^(***β***). When the eigenspaces of **G** and **P** are more closely aligned, directional heritability tends to vary less across phenotypic directions, whereas misalignment tends to broaden the distribution and increase the mass of low-heritability directions. Changes in the strength and patterning of covariances in **G** can also affect the spread of *h*^2^(***β***), but these effects enter the distribution primarily through how they reshape **G**^∗^ on the phenotypic scale. Figure 3 summarises several qualitative regimes produced by different combinations of spectra and alignment, and Figure 4 provides a concrete two-trait illustration of the same mechanism: misalignment expands the range of *h*^2^ across directions and creates low-heritability regions even when genetic variance is not uniformly small.

**Figure 3.**
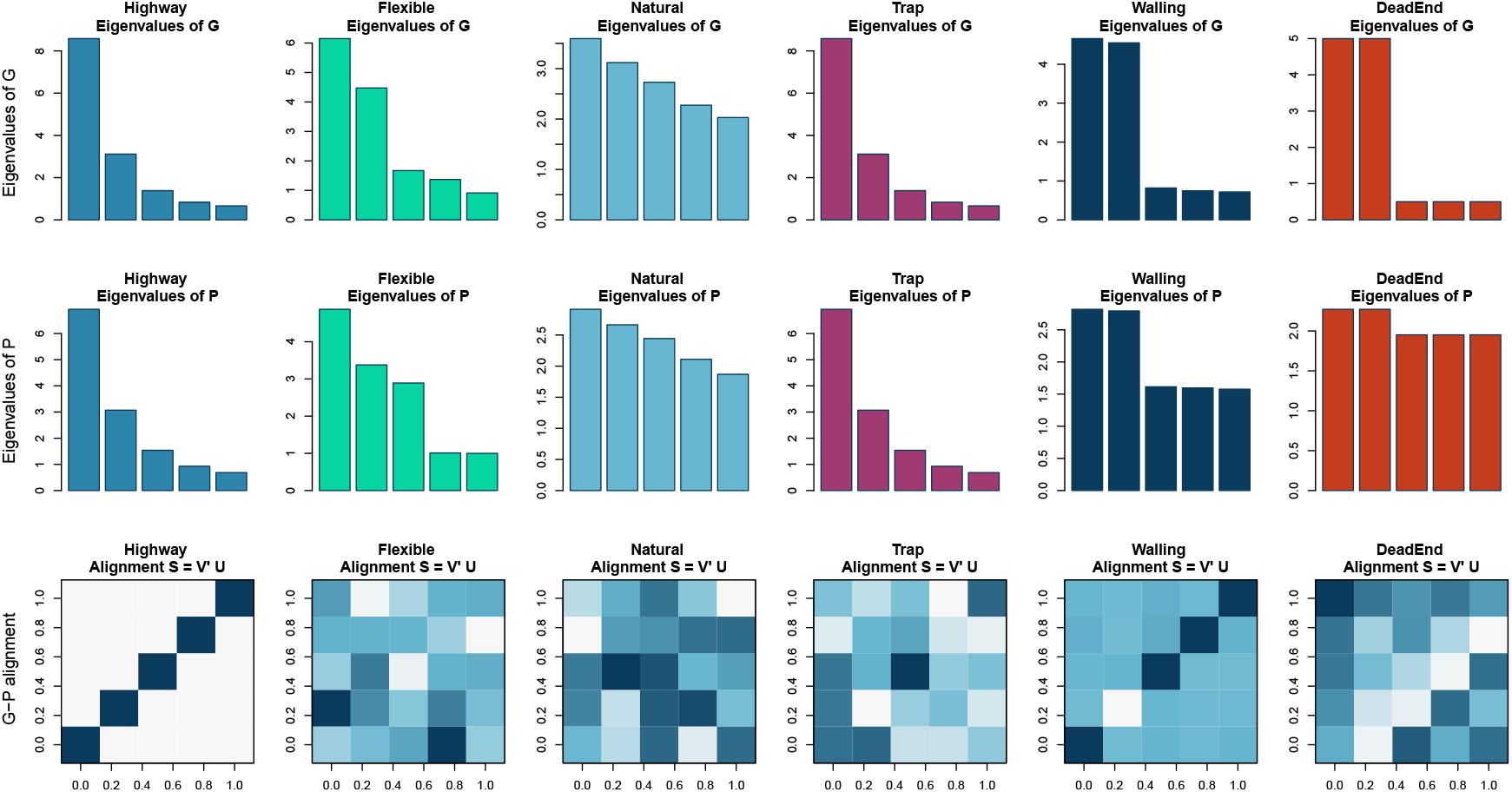
Eigenvalue structure and G-P alignment across six evolutionary scenarios. Each column represents a biologically interpretable scenario ranging from favourable (Highway) to severely constrained (DeadEnd) evolutionary conditions. *Top row*: Eigenvalue spectra of **G** showing genetic variance distribution across principal axes. *Middle row*: Eigenvalue spectra of **P** showing phenotypic variance distribution. *Bottom row*: Alignment matrices **S** = **V**^*′*^**U**, where **U** and **V** are eigenvector matrices of **G** and **P** respectively. Dark diagonal elements indicate aligned axes; off-diagonal darkness indicates misalignment. **Highway**: Strong correlations with perfect G-P alignment create an evolutionary “highway” along the leading eigenvector. **Flexible**: Weak correlations permit evolution in multiple directions despite misalignment. **Natural**: Intermediate alignment and correlation typical of natural populations. **Trap**: Strong correlations with misalignment create evolutionary “traps” where genetic variance points away from phenotypically variable directions. **Walling**: High eigenvalue dispersion with near-perfect alignment, analogous to agricultural breeding programmes. **DeadEnd**: Severe misalignment combined with clustered eigenvalues creates the most constrained evolutionary scenario.

**Figure 4.**
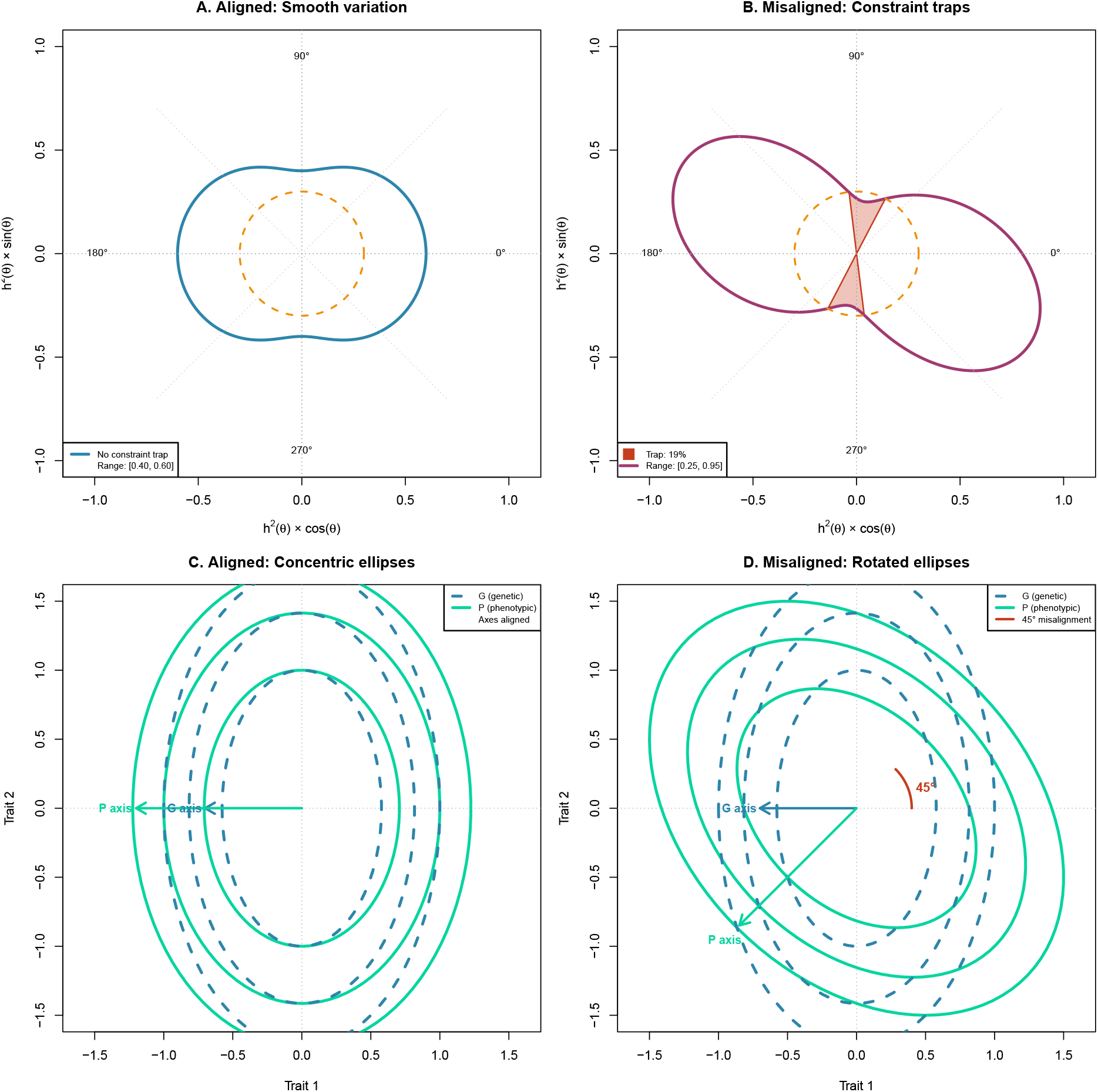
Directional heritability and constraint traps in two-trait systems. Two-dimensional examples illustrating how G-P misalignment creates constraint traps. *Top row*: Polar plots of directional heritability *h*^2^(*θ*) as a function of selection direction *θ*. Distance from origin indicates *h*^2^ magnitude; the dashed circle marks the constraint threshold (*h*^2^ = 0.3). **(A)** Aligned case: *h*^2^(*θ*) varies smoothly between generalized eigenvalues *λ*_min_ = 0.40 and *λ*_max_ = 0.60, with all directions above threshold. **(B)** Misaligned case (45°rotation): *h*_2_(*θ*) range expands to [0.25, 0.95], creating constraint traps (shaded red regions) comprising 19% of selection directions where *h*^2^ *<* 0.3. *Bottom row*: Variance ellipses showing genetic (**G**, dashed blue) and phenotypic (**P**, solid teal) covariance structure. **(C)** Aligned: G and P ellipses share principal axes, yielding smooth *h*^2^(*θ*) variation. **(D)** Misaligned: 45°rotation between G and P axes creates directions where genetic variance is low but phenotypic variance is high (constraint traps) or vice versa.

These simulations are deliberately stylised. They are intended to clarify how the distributional summary depends on matrix geometry under the uniform-on-**P**-sphere baseline, not to reproduce the details of any particular empirical system or any particular realised distribution of selection gradients.

### Empirical Application

#### Data

We compiled 55 **G**–**P** matrix pairs from 11 published studies spanning birds, mammals, fish, lizards, insects, and plants (Table 2). Matrix dimensions ranged from 2 to 8 traits. For each pair, we sampled 10,000 random selection directions on the **P**-unit sphere and computed directional heritability for each. All data analysis was conducted in R 4.5.2 (R Core Team 2025).

**Table 2.**
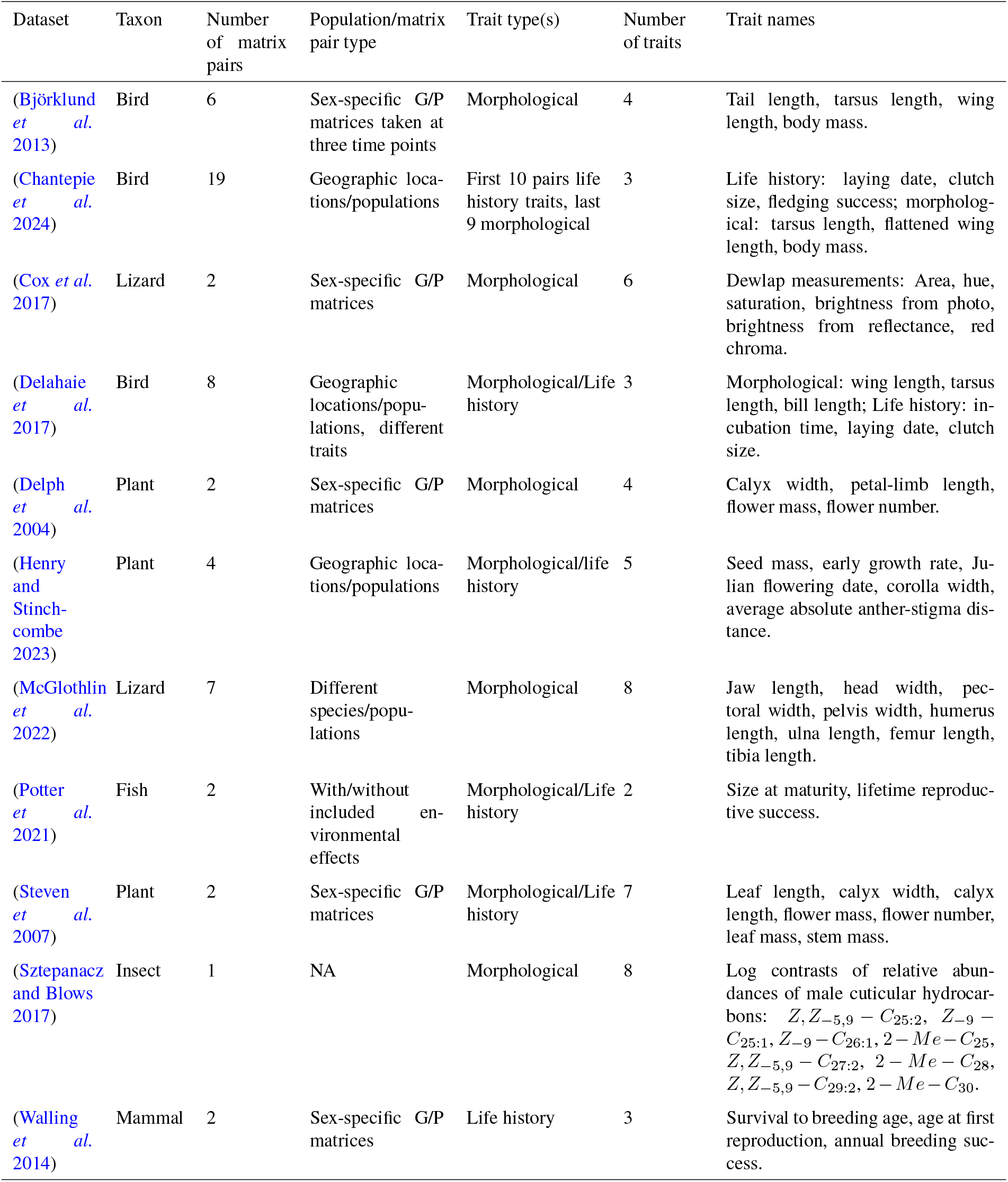
Empirical datasets and features of their **G** and **P** matrices.

| Dataset | Taxon | Number of matrix pairs | Population/matrix pair type | Trait type(s) | Number of traits | Trait names |
| --- | --- | --- | --- | --- | --- | --- |
| (Björklund *et al.* 2013) | Bird | 6 | Sex-specific G/P matrices taken at three time points | Morphological | 4 | Tail length, tarsus length, wing length, body mass. |
| (Chantepie *et al.* 2024) | Bird | 19 | Geographic locations/populations | First 10 pairs life history traits, last 9 morphological | 3 | Life history: laying date, clutch size, fledging success; morphological: tarsus length, flattened wing length, body mass. |
| (Cox *et al.* 2017) | Lizard | 2 | Sex-specific G/P matrices | Morphological | 6 | Dewlap measurements: Area, hue, saturation, brightness from photo, brightness from reflectance, red chroma. |
| (Delahaie *et al.* 2017) | Bird | 8 | Geographic locations/populations, different traits | Morphological/Life history | 3 | Morphological: wing length, tarsus length, bill length; Life history: incubation time, laying date, clutch size. |
| (Delph *et al.* 2004) | Plant | 2 | Sex-specific G/P matrices | Morphological | 4 | Calyx width, petal-limb length, flower mass, flower number. |
| (Henry and Stinchcombe 2023) | Plant | 4 | Geographic locations/populations | Morphological/life history | 5 | Seed mass, early growth rate, Julian flowering date, corolla width, average absolute anther-stigma distance. |
| (McGlothlin *et al.* 2022) | Lizard | 7 | Different species/populations | Morphological | 8 | Jaw length, head width, pectoral width, pelvis width, humerus length, ulna length, femur length, tibia length. |
| (Potter *et al.* 2021) | Fish | 2 | With/without included environmental effects | Morphological/Life history | 2 | Size at maturity, lifetime reproductive success. |
| (Steven *et al.* 2007) | Plant | 2 | Sex-specific G/P matrices | Morphological/Life history | 7 | Leaf length, calyx width, calyx length, flower mass, flower number, leaf mass, stem mass. |
| (Sztepanacz and Blows 2017) | Insect | 1 | NA | Morphological | 8 | Log contrasts of relative abundances of male cuticular hydrocarbons: $Z, Z_{-5,9} - C_{25:2}$, $Z_{-9} - C_{25:1}$, $Z_{-9} - C_{26:1}$, $2 - Me - C_{25}$, $Z, Z_{-5,9} - C_{27:2}$, $2 - Me - C_{28}$, $Z, Z_{-5,9} - C_{29:2}$, $2 - Me - C_{30}$. |
| (Walling *et al.* 2014) | Mammal | 2 | Sex-specific G/P matrices | Life history | 3 | Survival to breeding age, age at first reproduction, annual breeding success. |

#### Variation in constraint patterns

The distributions of directional heritability varied substantially across systems (Figure 5). The coefficient of variation ranged from 0.11 to 0.67, indicating that some populations experience relatively uniform heritability across selection gradients while others face heterogeneous constraint landscapes (Table S1). Mean directional heritability ranged from 0.02 to 0.53.

**Figure 5.**
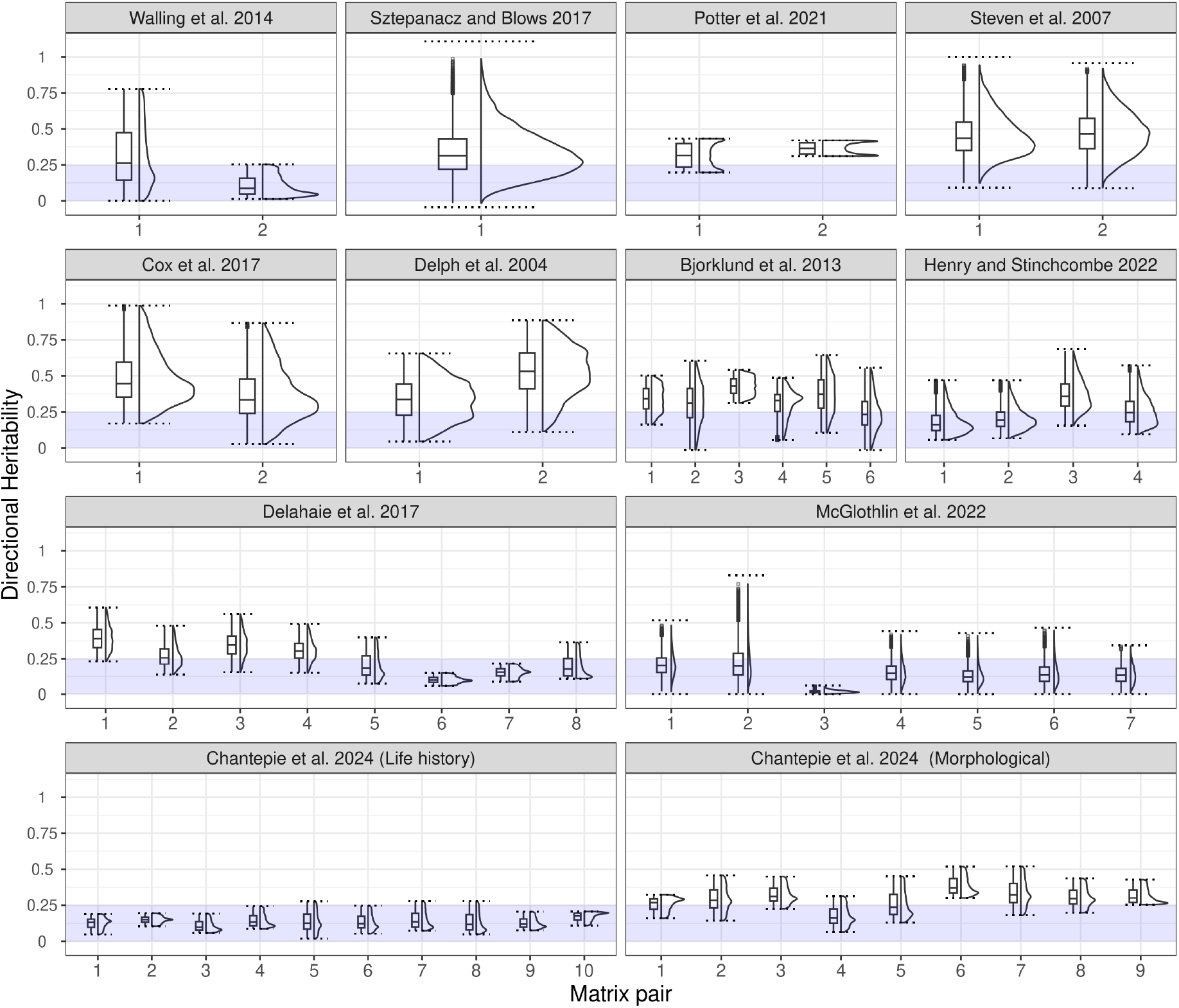
Empirical patterns in directional heritability. Distributions of *h*^2^(***β***) across 10,000 random directions for matrix pairs from multiple studies, showing variation in mean, spread, and skewness. Blue shading indicates constraint region (*h*^2^ *<* 0.25).

Constraint probabilities also varied widely. At a threshold of *h*^2^ = 0.25, the fraction of directions falling below this value ranged from 0% to over 90% across matrix pairs. 67.3% of populations examined had more than 25% of directions in this low-heritability region.

#### Misalignment and CV(*h*^2^)

To identify the association between **G** and **P** misalignment and variation in directional heritability, we ran a robust linear model (with HC2 errors) of the form Maximum misalignment ~ CV(*h*^2^(***β***)) across the matrix pairs where alignment could be reliably estimated. Misalignment between **G** and **P** eigenvectors was positively associated with CV(*h*^2^), with a 0.1-unit increase in CV(*h*^2^(***β***)) predicting a 14.8^°^ increase in maximum misalignment (Figure 6; *F*_1,53_ = 53.27, adjusted *R*^2^ = 0.38, *p <* 0.001). This pattern is consistent with the simulation results and suggests that the relative orientation of genetic and phenotypic variance structures contributes to variation in constraint heterogeneity across natural populations. Together, these summaries provide a compact description of how often and how strongly evolutionary responses are expected to be limited in each system.

**Figure 6.**
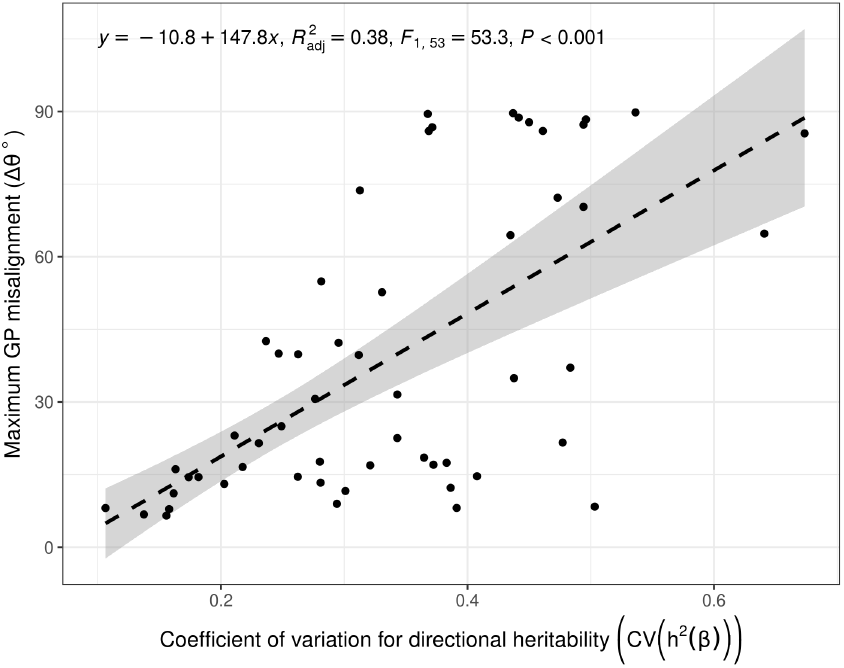
Relationship between G–P misalignment and CV(*h*^2^). Maximum canonical angle between eigenvector systems plotted against coefficient of variation of directional heritability for 55 matrix pairs from 11 studies. Each point represents a single matrix pair. Dashed line shows robust linear regression of misalignment ∼ *CV* (*h*^2^(***β***) with 95% confidence band (adjusted *R*_2_ = 0.38, *F*_1,53_ = 53.27, *p <* 0.001). Confidence band calculated via HC2. Equation in the top left is the model fit.

### Evolvability and Directional Heritability

#### Two measures, two determinants

Evolvability *e*(***β***) = ***β***^⊤^**G*β*** and directional heritability *h*^2^(***β***) address different aspects of evolutionary constraint. Evolvability measures the absolute genetic variance available in a direction; directional heritability measures the fraction of what selection perceives that is actually heritable. The breeder’s equation 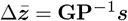 shows that both matter: selection acts on phenotypic differences, but only the genetic component transmits to offspring.

This distinction is reflected in their distributional properties. Under uniformly distributed selection directions, the spreads of these quantities have parallel forms but different determinants:

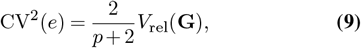

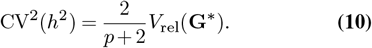

The spread of evolvability depends on **G** alone. The spread of directional heritability depends on **G**^∗^ = **P**^−1/2^**GP**^−1/2^, which absorbs the eigenvalue structures of both **G** and **P** together with their alignment. Consequently, CV(*h*^2^) responds to **G**–**P** alignment while CV(*e*) does not.

This asymmetry has practical consequences. Our simulations show that CV(*h*^2^) decreases roughly twofold as alignment increases from poor to good, whereas CV(*e*) remains nearly constant. Two populations with identical **G** matrices but different environmental variance structures will share the same evolvability distribution yet may have very different heritability distributions. Populations with misaligned **G** and **P** may face more constraint traps and greater evolutionary unpredictability – patterns invisible to evolvability analysis alone.

#### Empirical decoupling and three regimes

Figure 7 confirms that evolvability and directional heritability are largely uncorrelated across directions within populations, as previously observed (Houle 1992). Knowing the evolvability of a direction provides limited information about its heritability. This decoupling reveals three qualitatively distinct regimes in the joint distribution.

**Figure 7.**
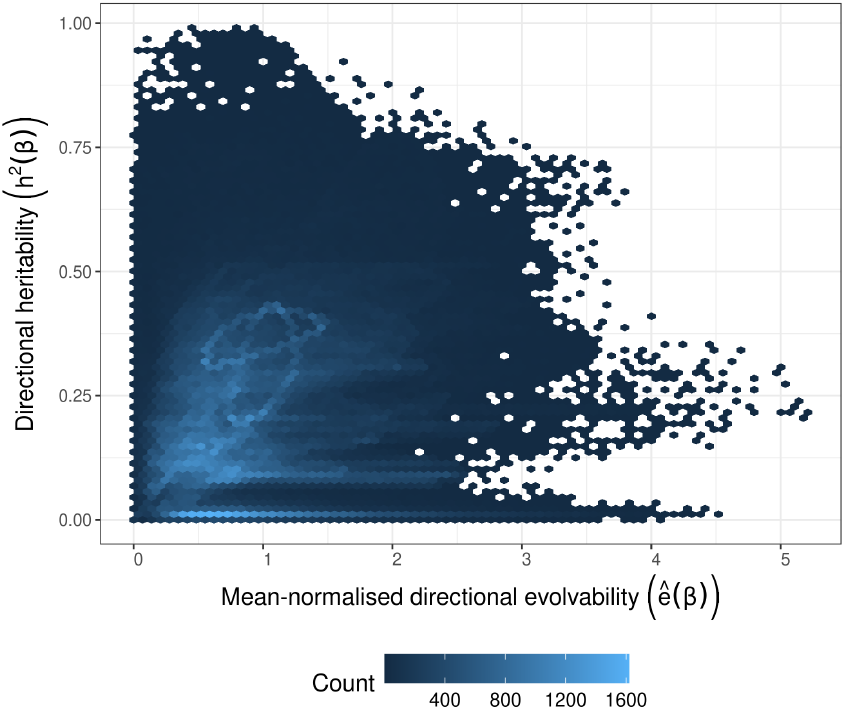
Relationship between directional heritability. (*h*^2^(***β***)) **and mean-normalised evolvability** (*ê*(***β***) = *e*(***β***)*/ē*(***β***) and *e*(***β***) = ***β***^⊤^**G*β***) **across 55 pairs of G and P matrices from 11 published studies**. Hex bins show counts of *h*^2^(***β***) and *ê*(***β***) outcomes across 10,000 randomly sampled ***β*** directions per matrix pair. *ē*(***β***) was measured across the 10,000 ***β*** directions per matrix pair. The two elliptical shapes in light blue around *ê*(***β***), *h*^2^(***β***) ≈ (1, 0.3) are from the Potter *et al*. (2021) dataset. The elliptical shape arises because this dataset contains two traits.

First, directions with high evolvability and high heritability represent favourable evolutionary opportunities – the open roads of phenotype space where evolution proceeds readily. Second, directions with low evolvability and low heritability are constrained in the classical sense: genetic variance is simply absent. Third, and most consequential for applied work, directions with moderate-to-high evolvability but low heritability constitute constraint traps. In these directions, genetic variance exists, but environmental variance is proportionally larger. Selection perceives phenotypic variation yet cannot efficiently extract the genetic component. A researcher pursuing such a direction based solely on evolvability would anticipate a reasonable evolutionary response but would observe weak genetic gain. How common this this response is unclear, but some examples of environmental variance hindering adaptive responses have been observed (see Alatalo *et al*. 1990).

We observed an elliptical pattern in the relationship between heritability and evolvability, centered at *h*^2^(***β***) ≈ 0.3 and *e*(***β***) ≈ 1 (Fig. 7). Upon further investigation, we found that these patterns were from the Potter *et al*. (2021) dataset. This dataset produced 2×2 matrices, explaining the elliptical shape. Potter *et al*. (2021) estimated **G** and **P** matrices with and without spatiotemporal subgroup effects (population density and breeding season). The two specifications yield visibly different evolvability/directional heritability relationships. In the **GP** pair without the subgroup effects, the distribution of *ê*(***β***) is wider, with smaller variation in *h*^2^(***β***). The opposite is true for the matrix pair including subgroup effects (Fig. 7). Including environmental cofactors in the estimates of **G** and **P** in this case reduces the spread of directional heritability, but increases evolvability, reflecting an “escape” from a constraint trap (Fig. 5). Researchers should be wary of the effects of un-modeled environmental covariance on inflating both **G** and **P**, as both *ê*(***β***) and *h*^2^(***β***) are dependent on the quality of matrix estimates.

#### Implications for prediction and breeding

The existence of constraint traps has direct implications for evolutionary prediction. Consider two populations: one where **G** and **P** share similar eigenvector orientations and another where they diverge. In the aligned population, directions of high genetic variance coincide with directions of high phenotypic variance. In this case, heritability is relatively stable across directions, and evolutionary trajectories are predictable from **G**-matrix geometry. In the misaligned population, some directions offer efficient evolutionary pathways while others are constraint traps. Evolutionary outcomes depend sensitively on where selection happens to push, and predictions based on **G** alone may mislead.

For breeding programmes, the distinction translates directly into the difference between genetic variance and response to selection. A breeding target in a constraint trap will show smaller-than-expected gains despite strong selection pressure. The phenotypic variation observed in the breeding population suggests potential for improvement, but most of that variation is environmental and does not transmit to offspring. By computing *h*^2^(***β***) for a proposed selection index, breeders can assess whether the target direction offers efficient genetic gain or falls into a constraint trap. If the latter, the index can be revised to steer toward more favourable directions.

More broadly, the constraint probability CR(*c*) measures how forgiving a genetic architecture is by capturing the fraction of possible selection targets that would encounter heritability below some acceptable threshold. Architectures with low CR(*c*) offer flexibility: selection can be applied in many directions with reasonable efficiency. Architectures with high CR(*c*) require careful choice of breeding objectives to avoid low-response directions.

## Discussion

How much does heritability vary across selection directions, and what determines this variation? We have shown that the answer depends on a single matrix: the **P**-whitened genetic variance matrix **G**^∗^ = **P**^−1/2^**GP**^−1/2^. The eigenvalue dispersion of this matrix, combined with the number of traits, fully determines the coefficient of variation of directional heritability under uniformly distributed selection on the **P**-unit sphere. This result reduces the problem of characterising multivariate constraint from comparing two matrices to examining one.

Our choice of baseline – uniform directions on the **P**-sphere – is not intended as a literal model of how selection is generated in nature. It is a reference that makes “random phenotypic directions” well-defined and comparable across trait scalings and across studies, because directions are measured in units of phenotypic variance rather than in arbitrary Euclidean coordinates. When empirical information about the distribution of selection gradients is available, the same calculations can be carried out under that distribution; our results then provide a baseline against which structured selection can be compared.

### Geometry, alignment, and constraint

Perhaps the most unexpected finding from our simulations is that alignment between the eigenspaces of **G** and **P** can matter more than the magnitude of genetic correlations. This result deserves reflection. Evolutionary geneticists have long focused on correlation structure as the primary determinant of constraint, and with good reason: strong genetic correlations channel evolutionary trajectories along particular paths (Schluter 1996). Yet our analysis suggests that the relative orientation of genetic and phenotypic variance may be equally consequential.

The intuition is geometric. When **G** and **P** share similar principal axes, genetic variance tends to be concentrated in directions where phenotypic variance is also high. The whitening transformation then compresses rather than stretches the eigen-value spectrum, keeping CV(*h*^2^) modest even when raw eigen-value ratios in **G** are extreme. This may help explain empirical patterns, such as those in Walling *et al*. (2014), where populations with strong genetic correlations nevertheless show relatively stable heritability across directions.

Conversely, misalignment can inflate *V*_rel_(**G**^∗^) even when both **G** and **P** individually have moderate eigenvalue dispersion. The biological consequence is that some phenotypic directions (potentially those favoured by novel selection) may harbour substantial variance yet offer little heritable signal. These constraint traps are invisible to analyses focused on **G** alone.

We suspect that the relative importance of alignment versus correlation strength varies across biological systems in ways that remain to be characterised. Populations experiencing stable environmental conditions may maintain alignment through consistent selection, while those in fluctuating environments may accumulate misalignment as environmental variance shifts independently of genetic architecture (Vinton *et al*. 2022). The presence of substantial environmental variance could hinder the ability of selection to target genetic variance, reducing the response to selection despite substantial genetic variance. This has large implications for the evolution of phenotypic plasticity. For instance, when selection acts along a constraint trap, and there is substantial environmental variance, plasticity should be selected for. Following this, genetic assimilation might solidify the response in the genetic architecture, realigning genetic variance towards the direction of selection. Testing these hypotheses will require datasets where both matrices are estimated with sufficient precision to distinguish true misalignment from sampling error.

### Future work

Three observations emerge from this work that may warrant further investigation.

First, the whitened matrix **G**^∗^ serves as a sufficient summary for directional heritability variation under the baseline model of uniformly distributed selection on the **P**-sphere. Its eigenvalue dispersion, combined with trait dimensionality, determines how much heritability varies across selection directions under that model. The relationship CV^2^(*h*^2^) = (2/(*p* + 2))*V*_rel_(**G**^∗^) is exact under the stated assumptions. Whether the simplification from two matrices to one proves useful in practice will depend on how well the baseline approximates realised selective regimes.

Second, empirical systems span a broad range of constraint heterogeneity. Across the matrix pairs we examined, the coefficient of variation of directional heritability ranged from 0.11 to 0.67. Some populations exhibit relatively uniform heritability regardless of selection direction; others face landscapes in which heritability varies several-fold. We do not yet know what drives this variation across taxa and trait types, though our framework suggests that **G**–**P** alignment is one contributing factor. Understanding the ecological and evolutionary determinants of constraint heterogeneity seems a promising direction for future work.

Third, evolvability and directional heritability provide complementary rather than redundant information (Figure 7). The lack of correlation between these quantities across directions within populations means that knowing one tells us little about the other (Houle 1992). This decoupling is not a failure of either measure but a reflection of biological reality: genetic variance and the ratio of genetic to phenotypic variance are simply different properties of a genetic architecture. Directional heritability is most useful as a phenotypic-scale efficiency diagnostic, particularly in directions where phenotypic variance is substantial and the transmissible fraction may be small. In such settings, *h*^2^(***β***) can identify directions where phenotypic variation masks a heritable signal even if genetic variance is present.

For the joint *e*–*h*^2^ diagnostic (Figure 7), we evaluate both quantities for the same set of candidate directions in the original trait space (including observed or proposed selection gradients or indices), rather than restricting directions to the **P**-unit sphere used for the distributional null. The distributional theory concerns an invariant baseline over phenotypic directions, whereas the joint plot is intended as a practical check on where particular directions lie relative to the architecture.

### Connections to prior work

Our analysis extends rather than replaces Hansen and Houle’s (2008) evolvability framework. Their focus on average evolvability and conditional evolvability provided tools for summarising **G** that have proven valuable across many empirical systems. We maintain their geometric interpretation but shift attention from averages to distributions, and from **G** alone to the **G**–**P** relationship captured by the whitened **G**^∗^ matrix.

Directional heritability has a direct antecedent in selection index theory, where the heritability of an index is defined as a ratio of quadratic forms (Lin and Allaire 1977). Selection index theory typically treats this quantity as a property of a chosen index; our contribution is to treat the same quantity as a landscape over directions and to quantify its heterogeneity and tail risk across direction space. The distributional viewpoint also connects to recent work treating evolutionary quantities as quadratic forms or ratios of quadratic forms evaluated on random directions (Watanabe 2024). Our results identify a transformation that isolates the relevant object for directional heritability: after **P**-whitening, *h*^2^(***β***) becomes a single quadratic form in **G**^∗^ evaluated on a Euclidean unit vector, clarifying why **G**–**P** alignment enters strongly into constraint heterogeneity.

Finally, null models in quantitative genetics have long used eigenvalue spectra to distinguish structured genetic organisation from unstructured baselines (Wagner 1984). Our framework is compatible with this tradition: once a null for the joint structure of (**G, P**) is specified, it implies corresponding expectations for the generalized eigenvalues and thus for the spread and tail probabilities of *h*^2^(***β***). In that sense, CV(*h*^2^) and CR(*c*) provide compact summaries that can be compared to null expectations and used to diagnose non-random organisation of transmissible signal across phenotype space.

### Limitations and open questions

Several limitations should temper interpretation of our results.

Our theoretical framework assumes that selection directions are uniformly distributed on the **P**-unit sphere. Real selection is unlikely to be uniform; some directions may be targeted repeatedly while others are never encountered. In a meta-analysis of 60 studies of five populations, Siepielski *et al*. (2013) found that variability in the direction of selection was weaker than variation in the strength of selection, supporting this hypothesis. If selection is concentrated along axes that happen to have high (or low) heritability, the distribution of realised evolutionary responses would differ substantially from our predictions. Incorporating empirical distributions of selection gradients (perhaps from databases of measured selection differentials) would extend the framework toward more realistic scenarios.

Sampling error in estimated **G** and **P** matrices propagates to all derived quantities. Given the large sample sizes required to estimate **G**, this error can be substantial (Arnold *et al*. 2008). The eigenvalue ratios exceeding biologically plausible bounds in some of our empirical matrices almost certainly reflect estimation error rather than biological reality. Developing methods to propagate uncertainty through the whitening transformation and into distributional summaries would strengthen empirical applications of this framework. Our analyses span matrices with 2–8 traits, but multivariate constraint in nature involves many more dimensions. Whether the patterns we observe scale to higher-dimensional systems, or whether effective dimensionality saturates as trait number increases, remains unclear.

The framework also inherits assumptions from the underlying estimation procedures. Most empirical **G**-matrices are estimated from breeding designs or animal models that assume multivariate normality. Departures from normality, particularly in the environmental component, could affect the distribution of directional heritability in ways our Gaussian-based derivations do not capture. Another problem is that estimates of additive variance (i.e. the **G** matrix) can be inflated by the presence of genotype-environment interactions (Munar-Delgado *et al*. 2023). Furthermore, **G** and **P** are often estimated from different data sources or with different sample sizes, introducing covariance between estimation errors that complicates uncertainty propagation. As trait dimensionality increases, these estimation challenges compound: the number of parameters in **G** and **P** grows quadratically while information per parameter typically shrinks.

We have treated **G** and **P** as fixed, but both matrices evolve (Arnold *et al*. 2008). Strong directional selection depletes genetic variance along the selected axis (Barton and Turelli 1989), although fluctuating selection may maintain variance more broadly (Sasaki and Ellner 1997). Environmental change alters **P** directly. How the distribution of directional heritability shifts as populations adapt — and whether populations can evolve toward architectures with lower constraint risk — are questions our static framework cannot address.

Finally, our empirical analyses draw on matrix pairs from published studies, each with its own estimation methods, sample sizes, and trait choices. The variation we observe in constraint heterogeneity across systems is real, but we cannot confidently attribute it to biological rather than methodological differences without more standardised comparisons.

### Looking forward

For empirical studies of multivariate constraint, we suggest that reporting both evolvability and heritability distributions may provide a more complete picture than either alone. Useful summaries include mean values (*ē* and 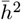), distributional spreads (CV(*e*) and CV(*h*^2^)), and constraint probabilities at biologically meaningful thresholds. The joint distribution of *e*(***β***) and *h*^2^(***β***) across candidate directions can reveal whether low-efficiency directions are common in a system and whether evolvability and heritability provide redundant or complementary information. Making these distributional summaries accessible to empiricists and breeders will require computational tools that propagate uncertainty through the whitening transformation and output interpretable summaries. Software that takes raw variance component estimates and returns distributional diagnostics (e.g. CV(*h*^2^), constraint probabilities, and joint *e*–*h*^2^ plots) would lower barriers to applying this framework in practice.

Testing whether observed constraint patterns reflect biological organisation or chance will require null models. Random matrix approaches, such as Wagner (1984)’s random pleiotropy model, where **G** and **E** arise from independent developmental processes, generate predictable distributional properties for the generalised eigenvalues of (**G, P**). Empirical distributions that deviate from such nulls would indicate non-random organisation of the genotype-phenotype map. Developing formal statistical tests based on this logic would connect our descriptive framework to hypothesis-testing machinery.

More broadly, the framework developed here raises questions that our analysis cannot answer. What ecological or life-history factors predict **G**–**P** alignment? Do populations under fluctuating selection maintain lower constraint heterogeneity than those under stable selection? Can breeding programmes exploit geometric insights to avoid low-efficiency directions? Selection index theory already provides tools for choosing index weights (Lin and Allaire 1977); our distributional perspective suggests that breeders might additionally screen candidate indices for their position in the *h*^2^(***β***) landscape, avoiding directions near *λ*_min_ even if those directions align with breeding goals. Whether such geometric screening improves long-term genetic gain remains to be tested. These questions connect matrix geometry to evolutionary and applied outcomes in ways that seem worth pursuing.

## Data Availability

Code and data for the empirical simulations and analysis are available at https://github.com/nobrien97/geodirh2_2026. Empirical matrices were obtained from published sources cited in the text and described in detail at https://github.com/nobrien97/geodirh2_2026.

## Acknowledgments

We thank colleagues for discussions and researchers who have made their covariance matrices publicly available. We thank Jan Engelstädter, Mark Blows, Mark Cooper, Steve Chenoweth, Dolph Schluter, Katrina MacGuigan, and Simone Blomberg for fundamental discussions in evolutionary quantitative genetics and multivariate algebra. This work was supported by the Australian Research Council Centre of Excellence for Plant Success in Nature and Agriculture (CE200100015).

## Supplementary Online Materials

### Appendix S1: Geometric intuition for genetic variance and heritability

This appendix is written for readers who are comfortable with basic vectors and matrices, but who do not think of themselves as mathematicians or quantitative geneticists. The goal is to explain, in plain language, why objects such as ***β***^⊤^**G*β*** keep appearing, what it means to “whiten” by **P**, and why the matrix

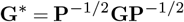

plays such a central role in the main text.

#### S1.1 Ellipses as pictures of variance

Think of a set of individuals measured for two traits, such as body size and flowering time. If you plot each individual’s traits as a point on a plane, the cloud of points usually looks like an elongated blob rather than a perfect circle.

Multivariate normal theory tells us that such a cloud can be summarised by:

- a mean vector (the centre of the cloud), and
- a variance–covariance matrix (the shape and orientation of the cloud).

For a 2 × 2 covariance matrix **M**, curves of equal probability density are ellipses:

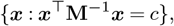

for different constants *c* > 0. The more stretched the ellipse, the more variance there is along that direction; the more tilted it is, the stronger the covariance.

In this geometric language:

- **G** is the ellipse describing how breeding values are spread in trait space;
- **P** is the ellipse describing how phenotypes are spread in trait space.

Everything that follows is really just different ways of reading information contained in these ellipses.

#### S1.2 Variance of a trait combination as a squared length

Suppose we combine *p* traits into a single index by taking a linear combination

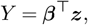

where ***z*** is the vector of trait values and ***β*** is a vector of weights (for example, regression coefficients for relative fitness). The variance of *Y* is

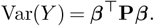

Algebraically, this is a *quadratic form*. Geometrically, it behaves like a squared length:

- in ordinary Euclidean space the squared length of a vector ***x*** is ***x***^⊤^**I*x***;
- in the “**P**-geometry” the squared length of ***β*** is ***β***^⊤^**P*β***.

So

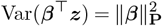

is “how long” the direction ***β*** is, when we measure length using the phenotypic variance structure as our ruler.

The same idea applies to additive genetic variance:

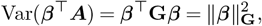

where ***A*** is the vector of breeding values. So **G** defines its own geometry on trait space: a direction can be long in the **G**-sense (lots of genetic variance) but not necessarily long in the **P**-sense (if phenotypic variance is small there), and vice versa.

#### S1.3 Directional heritability as a ratio of squared lengths

Directional heritability in direction ***β*** is

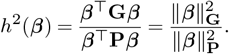

This is a ratio of two squared lengths:

- the numerator measures how much *genetic* variation lies along the direction ***β***;
- the denominator measures how much *phenotypic* variation lies along the same direction.

In other words, *h*^2^(***β***) is a signal-to-noise ratio: it tells us what fraction of the phenotypic variation that selection “sees” in direction ***β*** is transmissible to offspring.

This already explains why heritabilities can be low even when there is quite a lot of genetic variance: the denominator can be large because environmental variance happens to be large in that direction.

#### S1.4 Whitening: making phenotypic variance spherical

Working directly with the ratio

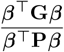

is awkward because both numerator and denominator change with ***β***. A standard trick in multivariate analysis is to transform coordinates so that one of the matrices becomes the identity. We do this with **P**.

Because **P** is symmetric and positive definite, it has a unique symmetric square root **P**^1/2^ and inverse square root **P**^−1/2^ satisfying

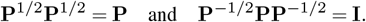

Define a new vector

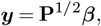

and then normalise it to unit Euclidean length:

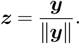

You can think of this as two steps:

1. **Change of units: *β*** ↦***y*** = **P**^1/2^***β*** rescales the axes so that **P** becomes the identity matrix. In this new coordinate system, phenotypic variance is the same in all directions: the **P**-ellipse has been turned into a circle.
2. **Normalisation:** we then force all directions to have Euclidean length 1 by dividing by ∥***y***∥.

In these whitened coordinates, the denominator of *h*^2^ simplifies:

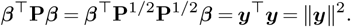

The numerator becomes

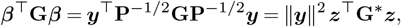

where

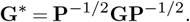

Putting numerator and denominator together, the factor ∥***y*** ∥^2^ cancels, and we obtain the simple expression

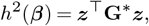

where ***z*** is a unit vector in ordinary Euclidean space.

This is the key simplification: *in whitened coordinates, directional heritability is just a quadratic form of a single matrix evaluated on a unit vector*. All the complexity of the ratio **G** vs. **P** has been absorbed into **G**^∗^.

#### S1.5 What the whitened matrix G^∗^ tells us

Because **G**^∗^ is symmetric, it has an eigendecomposition

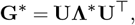

where **Λ**^∗^ is a diagonal matrix of real eigenvalues 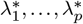 and **U** is an orthogonal matrix of eigenvectors.

Geometrically:

- the eigenvectors of **G**^∗^ are orthogonal directions in whitened trait space;
- the eigenvalues of **G**^∗^ are the heritabilities along these directions.

Indeed, if we take ***z*** to be one of the eigenvectors, say ***z*** = ***u***_*i*_, then

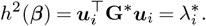

For a general unit vector ***z*** we can write ***z*** = **U*w***, where ***w*** is another unit vector consisting of the coordinates of ***z*** in the eigenbasis. Then

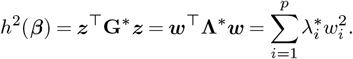

So directional heritability in whitened space is a weighted average of the eigenvalues of **G**^∗^, with non-negative weights 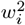 that sum to 1. When we sample directions at random (under the uniform-on-the-sphere assumption used in the main text), these weights have a very specific distribution, leading to the formula

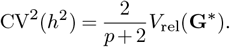

For the present appendix, the take-home messages are:

1. Any variance of a linear combination, such as ***β***^⊤^**P*β*** or ***β***^⊤^**G*β***, can be viewed as a squared length in the geometry induced by the corresponding matrix.
2. Directional heritability is a ratio of two such squared lengths, comparing the geometry of **G** to that of **P**.
3. Whitening by **P** turns this ratio into a single quadratic form of **G**^∗^, evaluated on unit vectors in ordinary Euclidean space.
4. The eigenvalues of **G**^∗^ are heritabilities along orthogonal directions in whitened space; their mean and spread summarise how much directional heritability varies across phenotypic directions.

Thinking geometrically in this way allows us to translate questions about “constraint” into questions about the shape of ellipses (or ellipsoids) and how stretched they are in different directions, which is often much easier to visualise than the underlying algebra.

##### S1.5.1 From geometry to distributions

The main text is not just about single directions, but about the *distribution* of quantities like evolvability and directional heritability across many possible selection directions.

Two points are especially important:

- Any variance in a given direction can be written as a quadratic form, such as *e*(***β***) = ***β***^⊤^**G*β*** or *h*^2^(***β***) = ***β***^⊤^**G*β***/***β***^⊤^**P*β***.
- After **P**-whitening, directional heritability becomes a single quadratic form of the whitened matrix,

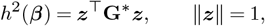

so the geometry of constraint is encoded by the eigenvalues and eigenvectors of **G**^∗^.

Once we see evolvability and heritability as *scalar fields on a sphere of directions*, we can ask distributional questions: if selection points in a random direction, what is the distribution of *h*^2^(***β***) or *e*(***β***) that it will encounter? Appendices 2 and 3 take this next step. Appendix S2 treats directional heritability *h*^2^(***β***) as a random variable and derives its mean and coefficient of variation in terms of the eigenvalues of **G**^∗^. Appendix S3 does the same for evolvability *e*(***β***) and shows how the two distributions are related but not redundant.

### Appendix S2: Directional heritability as a random variable

This appendix builds directly on Appendix S1. There we showed that, after **P**-whitening, directional heritability in direction ***β*** can be written as

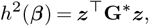

where ***z*** is a Euclidean unit vector (∥***z***∥ = 1) and

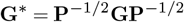

is the **P**-whitened genetic covariance matrix (Hansen and Houle 2008). The eigenvalues of **G**^∗^ are then heritabilities along orthogonal directions in whitened space.

Here we treat *h*^2^(***β***) as a *random variable* by allowing the direction of selection to wander randomly over the **P**-unit sphere. Our goal is to derive an expression for how much *h*^2^ varies across those directions:

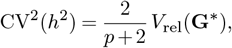

where *p* is the number of traits and *V*_rel_(**G**^∗^) is the relative eigenvalue variance of **G**^∗^,

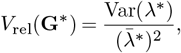

with 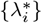 the eigenvalues of **G**^∗^ and 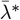 their mean (Pavlicev *et al*. 2009; Watanabe 2024). This result links the *distribution* of directional heritabilities across selection directions to the eigen-value spectrum of the whitened matrix.

#### S2.1 Step 1: Writing *h*^2^ as a weighted average of eigenvalues

Let the eigendecomposition of **G**^∗^ be

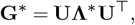

where 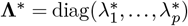 contains the eigenvalues and the columns of **U** are orthonormal eigenvectors.

Rotate the unit vector ***z*** into this eigenbasis:

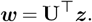

Because **U** is an orthogonal matrix, ***w*** is still a unit vector, so 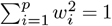.

Substituting into *h*^2^ gives

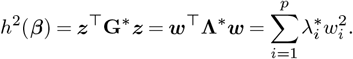

##### Key point

Directional heritability in a given direction is a weighted average of the eigenvalues of **G**^∗^, with non-negative weights 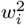 that sum to one. Geometry (the eigenvalues) now enters only through these weights.

#### S2.2 Step 2: Random directions and Dirichlet weights

In the main text we assume that selection directions are uniformly distributed on the **P**-unit sphere. In whitened coordinates, this means that ***z*** is uniformly distributed on the ordinary Euclidean unit sphere *S*^*p*−1^, and so is ***w*** = **U**^⊤^***z*** (a rotation preserves uniformity on the sphere).

A classical result from multivariate geometry is that when ***w*** is uniformly distributed on *S*^*p*−1^, the squared components

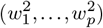

follow a symmetric Dirichlet distribution with all parameters equal to 1/2. The Dirichlet distribution describes how probability mass is allocated across categories; when the parameters are equal to 1/2, it corresponds to a uniform distribution on a sphere. We do not need the full machinery of this distribution here; only three simple facts about the moments:

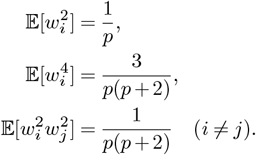

From these we obtain

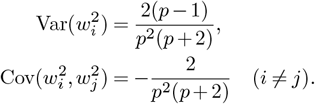

The negative covariance reflects a simple constraint: the weights must sum to one. If one 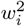 is unusually large, the others must be collectively smaller.

#### S2.3 Step 3: Mean directional heritability

Because *h*^2^ is a weighted sum of the eigenvalues,

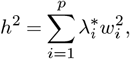

its expectation is

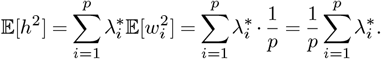

Thus

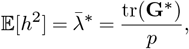

the mean eigenvalue of the whitened matrix. In words: the average directional heritability across random selection directions is simply the average of the heritabilities along the orthogonal axes defined by **G**^∗^.

#### S2.4 Step 4: Variance of directional heritability

To quantify the spread of directional heritabilities, we compute

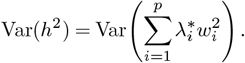

Expanding using the variance of a weighted sum,

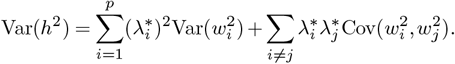

Substituting the expressions for 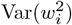 and 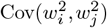 yields

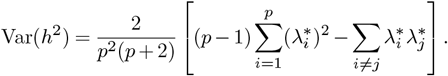

To simplify the bracketed term, write

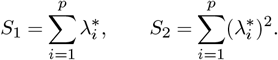

Then

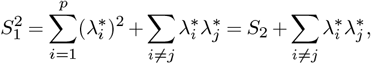

so

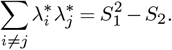

Substituting,

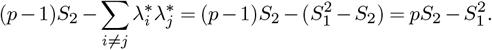

Hence

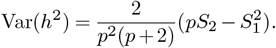

Now recall that the variance of the eigenvalues themselves is

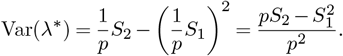

Combining these expressions gives the compact form

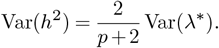

#### S2.5 Step 5: The CV^2^ formula and its meaning

The squared coefficient of variation of directional heritability is

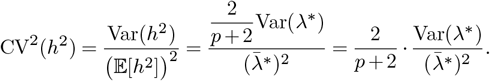

By definition,

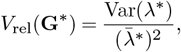

so we arrive at the main result:

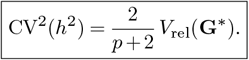

This expression makes the distributional message precise. Under uniformly distributed selection directions on the **P**-sphere:

- The *mean* directional heritability is set by the mean eigenvalue of 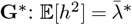.
- The *spread* of directional heritabilities around this mean is set by the relative eigenvalue variance of **G**^∗^, scaled by the factor 2/(*p* + 2) that comes from the geometry of random directions on a sphere.

In other words, the entire distribution of *h*^2^(***β***) across random selection directions is controlled by two simple ingredients: the dimensionality *p* and the eigenvalue dispersion of the whitened genetic matrix. This is the distributional link that underlies our empirical summaries of CV(*h*^2^) and constraint probabilities in the main text.

### Appendix S3: Distributions of evolvability and directional heritability

The main text emphasises that *distributions* of directional heritability and evolvability across directions are more informative than their averages. Appendix S2 derived the distributional spread of *h*^2^(***β***) under uniformly distributed selection directions on the **P**-sphere. Here we show that the same logic applies to evolvability *e*(***β***) = ***β***^⊤^**G*β***, and we clarify how the two distributions are related.

#### S3.1 Evolvability as a quadratic form on the Euclidean sphere

For evolvability, we do not have a ratio of quadratic forms. For any non-zero direction ***β*** in trait space,

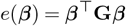

is already a single quadratic form in the Euclidean geometry defined by **G**. To put evolvability on the same footing as directional heritability, we again restrict attention to unit vectors and treat direction as the only degree of freedom:

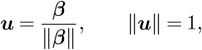

so that

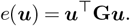

When selection directions are assumed to be uniformly distributed on the Euclidean unit sphere *S*^*p*−1^, evolvability becomes a random variable obtained by evaluating this quadratic form on random unit vectors.

#### S3.2 Parallels with Appendix S2

The algebra now proceeds exactly as in Appendix S2, but with two changes:

- We work with the genetic matrix **G** rather than the whitened matrix **G**^∗^.
- We draw directions uniformly from the Euclidean unit sphere (∥***u***∥ = 1) rather than from the **P**-sphere.

Write the eigendecomposition

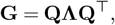

where **Λ** = diag(*λ*_1_, …, *λ*_*p*_) contains the eigenvalues of **G** and **Q** is orthogonal. Rotating into this eigenbasis, ***v*** = **Q**^⊤^***u***, we obtain

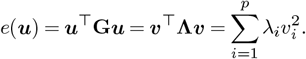

Thus evolvability in a given direction is a weighted average of the eigenvalues of **G**, with weights 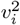 that are non-negative and sum to one. When ***u*** is uniformly distributed on the Euclidean sphere, 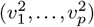 again follow a symmetric Dirichlet distribution with parameters 1/2, so

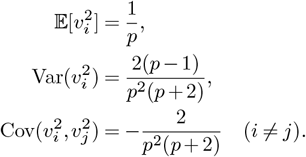

Repeating the same calculations as in Appendix S2, but with *λ*_*i*_ instead of 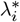, yields

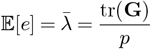

and

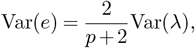

where Var(*λ*) is the variance of the eigenvalues of **G**. Defining

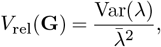

we arrive at the evolvability analogue of the main result:

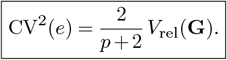

#### S3.3 Two distributions, two matrices

Taken together with Appendix S2, these results show that:

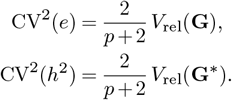

The two distributions are therefore controlled by different matrices:

- The distribution of *evolvabilities* across random directions depends on the eigenvalue spectrum of **G** itself. This spectrum reflects the anisotropy of additive genetic variance, shaped by mutation, selection and drift.
- The distribution of *directional heritabilities* across random directions depends on the eigenvalue spectrum of the whitened matrix **G**^∗^ = **P**^−1/2^**GP**^−1/2^. This spectrum folds in the geometry of phenotypic variance and the alignment between **G** and **P**.

The factor 2/(*p* + 2) appears in both expressions and comes from the same source: the geometry of random directions on a sphere, encoded by the Dirichlet distribution of squared components. The difference lies entirely in whether we are averaging the eigenvalues of **G** or those of **G**^∗^.

#### S3.4 Why the distributions are not redundant

Although the formulas for CV^2^(*e*) and CV^2^(*h*^2^) look similar, the underlying distributions carry different biological information:

- CV(*e*) tells us how patchy genetic variance is across directions *before* we consider phenotypic variance. High CV(*e*) means that some trait combinations have much more genetic variance than others.
- CV(*h*^2^) tells us how patchy the *fraction* of phenotypic variance that is genetic is across directions. High CV(*h*^2^) means that some directions offer efficient genetic response to selection while others are dominated by environmental noise.

Because **G**^∗^ depends on **G, P**, and their alignment, CV(*h*^2^) is sensitive to **G**–**P** misalignment whereas CV(*e*) is not. Our simulations show that:

- Increasing misalignment between **G** and **P** can substantially increase CV(*h*^2^), making the distribution of directional heritabilities more uneven.
- The same increase in misalignment leaves CV(*e*) largely unchanged, because the eigenvalues of **G** itself are unaffected.

Empirically (Figure 7), this difference shows up as a weak or absent correlation between *e*(***β***) and *h*^2^(***β***) across directions within populations. The joint distribution contains three regimes of interest:

1. High *e*, high *h*^2^: directions where genetic variance is abundant and efficiently expressed on the phenotypic scale.
2. Low *e*, low *h*^2^: genuinely constrained directions with little genetic fuel for evolution.
3. Moderate/high *e*, low *h*^2^: “constraint traps” where genetic variance exists but phenotypic variance is even larger, so selection sees noisy phenotypic differences and makes little genetic progress.

Constraint traps are common in our empirical datasets and are invisible if we look only at the distribution of *e*(***β***). They emerge only when we consider the distribution of *h*^2^(***β***) and its dependence on **G**^∗^.

In summary, Appendix S1 set up the geometric language for thinking about evolvability and directional heritability as quadratic forms. Appendix S2 showed that, under uniformly distributed selection directions on the **P**-sphere, the distribution of directional heritability *h*^2^(***β***) is controlled by the eigenvalue dispersion of the whitened matrix **G**^∗^. This appendix shows that, under uniformly distributed directions on the Euclidean sphere, the distribution of evolvability *e*(***β***) is controlled by the eigenvalue dispersion of **G** itself.

Together, these results justify our focus in the main text on:

- the mean and coefficient of variation of directional heritability and evolvability across directions;
- constraint probabilities such as CR(0.25), which read off tail mass from the *h*^2^ distribution;
- the joint distribution of *e*(***β***) and *h*^2^(***β***), which reveals constraint traps and clarifies how much evolutionary information is lost when we consider evolvability alone.

One take home message is that the eigenvalue spectra of **G** and **G**^∗^ provide compact summaries of the *distributions* of evolvability and directional heritability that selection is likely to encounter, and therefore of the multivariate constraints that shape evolutionary trajectories.

### Appendix S4: Tracking constraint under changing G and P

#### S4.1 Motivation

The main text treats **G** and **P** as fixed snapshots, which is appropriate when asking: given these estimated matrices, how is directional heritability distributed across phenotypic space? But both matrices change through time. Directional selection can deplete genetic variance along favoured axes (Barton and Turelli 1989); mutation and drift can replenish it elsewhere; environmental conditions can shift, altering the covariance structure of non-transmissible variation. A natural question follows: do the distributional tools we have developed, namely, *V*_rel_(**G**^∗^), CV(*h*^2^), and constraint probability CR(*c*), remain applicable when **G** and **P** are moving targets?

The main-text identity (Eq. 7) describes the distribution of *h*^2^(***β***) *conditional on* a given pair (**G, P**) under the uniform-on-**P**-sphere baseline. It is agnostic about how the matrices arose. If matrices change through time, the same identity applies at each time point *t*, conditional on (**G**_*t*_, **P**_*t*_), under the same baseline. The simulations below provide a numerical check of this snapshot-by-snapshot expectation, and a check that whitening and uniform direction sampling have been implemented consistently across a range of evolving matrix states.

#### S4.2 Definitions and two kinds of risk

At generation *t*, let **G**_*t*_ denote the additive genetic covariance matrix and **E**_*t*_ the covariance of non-transmissible effects. Phenotypic covariance is

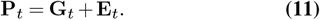

Directional heritability along selection gradient ***β*** is

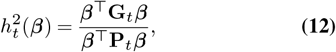

and the whitened genetic matrix is

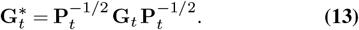

Under uniformly distributed selection directions on the **P**_*t*_-sphere,

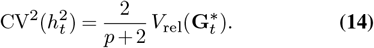

Operationally, we draw **z** uniformly from the Euclidean unit sphere *S*^*p*−1^ and compute 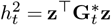. This is equivalent to drawing ***β*** uniformly on the **P**_*t*_-sphere via 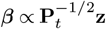.

When matrices evolve and selection direction may change, it is useful to distinguish two related but conceptually distinct questions. The first concerns *landscape risk*: at time *t*, what fraction of all possible phenotypic directions have low heritability? This is a property of the genetic architecture (**G**_*t*_, **P**_*t*_) at that moment, summarised by the constraint probability 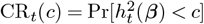, where the probability is taken over the **P**_*t*_-sphere baseline. Landscape risk describes the terrain—how much of phenotypic space consists of constraint traps.

The second concerns *realised risk*: at time *t*, does the particular direction that selection is actually pushing have low heritability? This is a single number, *h*^2^(***β***_*t*_), for the selection gradient applied at that generation. Realised risk describes the path—whether selection stepped into a trap.

A population can have low landscape risk yet experience high realised risk if selection targets one of the few constraint traps, or vice versa. A mountain where 25% of the surface is cliff illustrates the point: a population grazing on the plateau is safe; one whose path crosses cliff edges is not. Both perspectives are informative: landscape risk characterises architectural flexibility, realised risk characterises the actual trajectory.

#### S4.3 A phenomenological model of matrix change

We require a model that generates evolving **G**_*t*_ matrices without a full population-genetic simulation. The following update represents three generic contributions to change in standing variance:

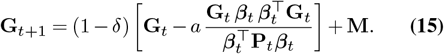

Each term has an interpretation. The subtracted term represents directional depletion—a heuristic reduction of variance in the selected direction, loosely motivated by the Bulmer effect (Bulmer 1971). We do not attempt a mechanistic linkage-disequilibrium treatment; the parameter *a* > 0 controls the rate at which variance erodes along ***β***_*t*_. The factor (1 ™ *δ*) represents global shrinkage from drift-like decay, with *δ ≥* 0 controlling uniform loss. The added matrix **M** represents mutational input of new variance each generation. After each update, we floor eigenvalues to maintain positive definiteness.

Environmental covariance **E**_*t*_ can also change. We model this by rotating the eigenvectors of **E**_*t*_ at a specified rate while holding eigenvalues constant, representing shifts in which trait combinations experience correlated environmental perturbations.

This model is deliberately stylised. Its purpose is to generate diverse (**G**_*t*_, **P**_*t*_) trajectories for numerical checking, not to serve as a realistic evolutionary simulator.

#### S4.4 Parameter sweep design

To check that the theoretical relationship holds across diverse matrix states, we conducted a parameter sweep varying initial heritability level (low, moderate, high), initial **G**–**P** misalignment (0°, 30°, 60°), rate of environmental covariance rotation, and rate of selection direction rotation. We simulated 108 parameter combinations, each run for 80 generations with snapshots every 10 generations, yielding 864 matrix states. At each snapshot, we computed theoretical predictions from the eigenvalues of **G**^∗^ and compared them to Monte Carlo estimates from 3,000–5,000 directions sampled uniformly on the **P**_*t*_-sphere.

#### S4.5 Results

Figure 2 provides the main numerical check. Each point is a snapshot *t* from the sweep, with 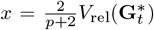 and 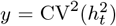 estimated by Monte Carlo sampling. Across 864 snapshots from 108 parameter combinations, the Monte Carlo estimates closely follow the theoretical relationship 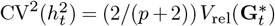 (descriptive *R*^2^ = 0.999, regression through the origin). The small departures from the line are consistent with finite Monte Carlo sampling and numerical tolerance. The colour scale indicates that this agreement holds across snapshots spanning mean directional heritabilities of roughly 0.15 to 0.55.

Figure S1 adds additional detail. Panel A shows that the mean relationship 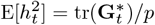 also matches the Monte Carlo estimates (descriptive *R*^2^ = 1.000). Panels B and C show that CR_*t*_(0.25) varies with both mean heritability and whitened anisotropy, which is consistent with both contributing to tail mass. However, CR_*t*_(*c*) depends on the full eigenvalue spectrum, not only its first two moments; mean and *V*_rel_ are informative summaries but they do not determine tail probabilities. Panel D shows that the same relationship in Figure 2 holds across initial misalignments (marker shapes) and heritability levels (colour).

Figure S2 illustrates the distinction between landscape and realised risk using two scenarios with identical initial conditions but different selection dynamics. In Panel A, selection remains fixed along the leading eigenvector of **G**_0_ (the initial genetic covariance matrix). Although the landscape risk starts high (CR ≈ 0.8) and averages 25% across the simulation, the realised heritability *h*^2^(***β***) stays above the threshold throughout. The population walks a safe corridor through moderately dangerous terrain. In Panel B, selection direction rotates through trait space. The landscape risk is similar (mean CR = 25%), but the realised heritability dips below the threshold 45% of the time (purple points). Same terrain, different path, different outcome.

This distinction has practical implications. A breeder selecting along a favourable direction may achieve consistent genetic gain even in a population with high landscape risk. Conversely, fluctuating selection can encounter constraint traps even when landscape risk is low, if it wanders into unfavourable regions.

#### S4.6 Summary

This appendix makes three points. First, the identity 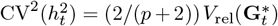 holds snapshot-by-snapshot under the uniform-on-**P**_*t*_-sphere baseline. The numerical check across 864 evolving matrix states (descriptive *R*^2^ = 0.999) is consistent with this expectation and provides a check on implementation.

Second, mean heritability and anisotropy contribute to constraint risk. Populations with low mean 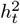 tend to face elevated constraint risk even at moderate 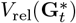, whereas populations with higher mean 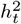 are buffered. However, CR_t_(*c*) depends on the full eigenvalue spectrum, so these summaries are informative but not sufficient.

Third, landscape and realised risk are distinct. A population can have dangerous terrain yet avoid traps by staying on a favourable path, or safe terrain yet encounter traps if selection wanders. Both metrics may be useful for understanding constraint in any particular system.

These results are numerical illustrations rather than predictions. A mechanistic treatment of **G**-matrix evolution under known mutation, selection, and drift would be needed for quantitative forecasts in specific systems. Our aim here is to show that the distributional tools introduced in the main text remain applicable—and that the theoretical relationships hold under the baseline—when the underlying matrices are not static.

**Figure S1.**
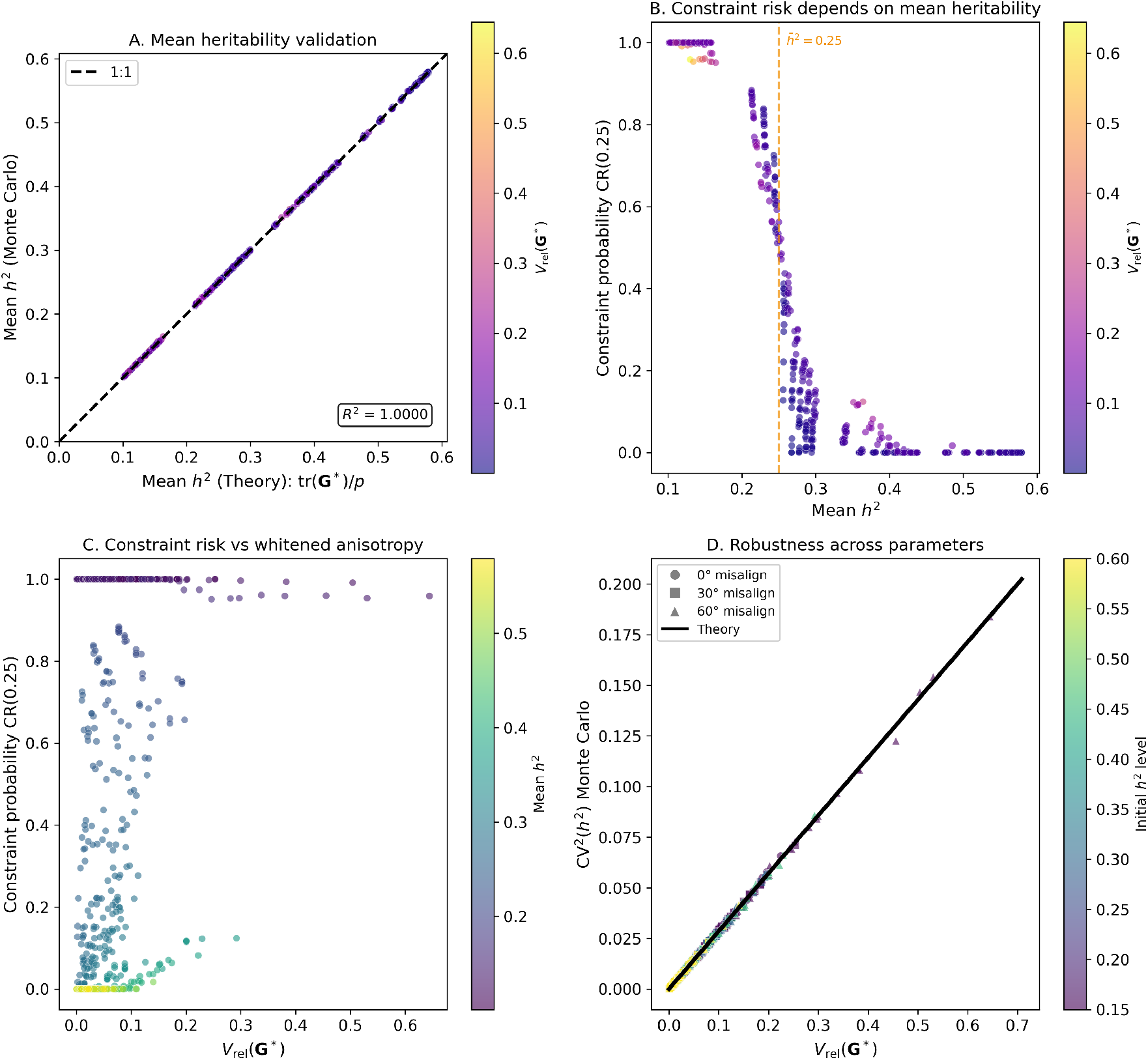
Robustness and contributors to constraint risk. (A) Mean heritability: theoretical prediction E[*h*^2^] = tr(**G**^∗^)*/p* versus Monte Carlo estimates (*R*^2^ = 1.000, descriptive). (B) Constraint probability CR(0.25) versus mean *h*^2^: low-heritability populations (purple) cluster at high constraint risk. (C) Constraint probability versus *V*_rel_(**G**^∗^): risk varies with both anisotropy and mean heritability (colour), though the full eigenvalue spectrum can also matter. (D) Parameter robustness: different initial misalignments (marker shapes) and heritability levels (colour) fall along the theoretical relationship.

### Appendix S5: Interactive companion application

#### S5.1 Purpose

To build geometric intuition for directional heritability, we provide an interactive Shiny application that allows users to manipulate two-trait systems and observe how eigenvalue structure and matrix alignment shape the joint distribution of evolvability and heritability across phenotypic directions. The app is pedagogical: it makes the whitening transformation, the distinction between Euclidean and **P**-sphere sampling, and the origin of elliptical patterns in (*e, h*^2^) space concrete and explorable.

#### S5.2 Availability

**Figure S2.**
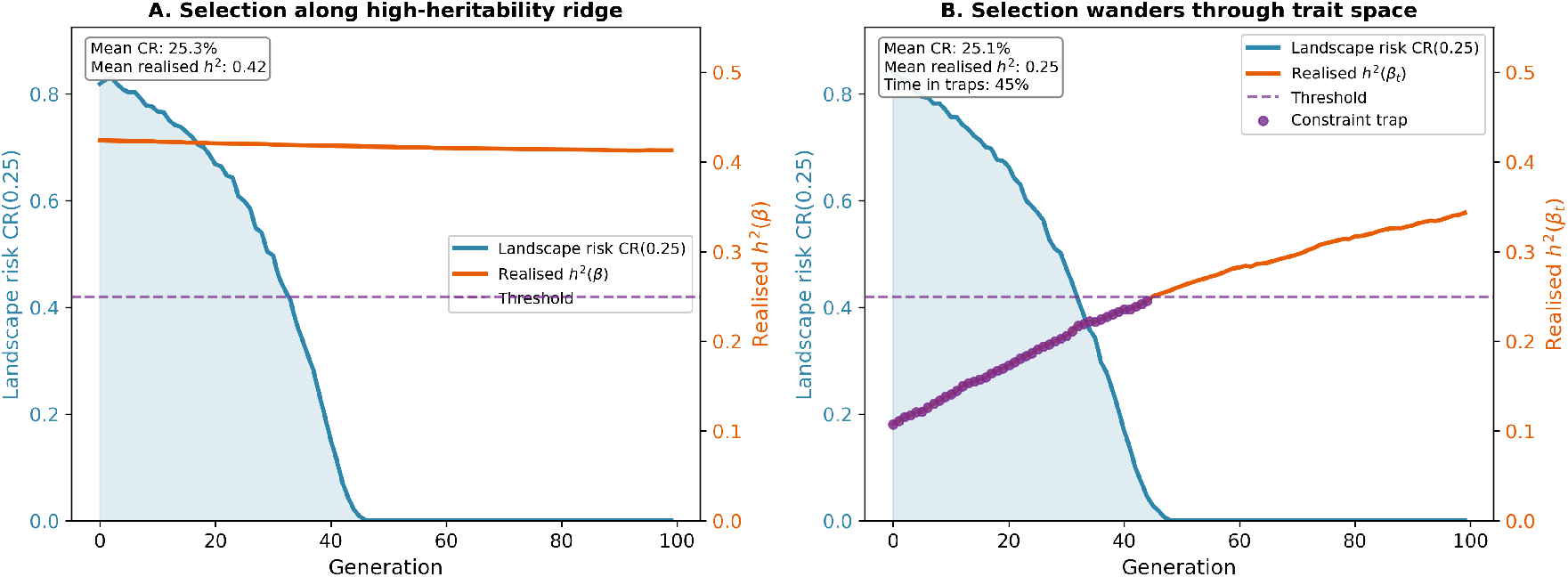
Landscape risk versus realised risk. Two scenarios with identical initial matrices but different selection dynamics. Blue shading: landscape risk CR(0.25); orange line: realised heritability *h*^2^(***β***_*t*_); purple dashed line: threshold *h*^2^ = 0.25; purple points: episodes where realised *h*^2^ falls below threshold. (A) Selection fixed along the leading eigenvector of **G**_0_: despite mean landscape risk of 25%, realised *h*^2^ stays above threshold throughout. (B) Selection rotates through trait space: similar landscape risk (25%), but realised *h*^2^ drops below threshold 45% of the time.

The application is available at https://dortizba.shinyapps.io/dirh2/ and can be run locally in R:

`library(shiny)`

`runApp(“app_dirh2.R”)`

The source code is available at https://github.com/nobrien97/geodirh2_2026.

#### S5.3 Interface overview

The application presents four panels. The upper-left panel shows the joint distribution of mean-normalised evolvability *e*(***β***) = ***β***′**G*β*** and directional heritability *h*^2^(***β***), with the user-specified constraint threshold *c* marked as a dashed line and the corresponding constraint probability CR(*c*) displayed. The upper-right panel shows the parametric (*e, p*) plot, where *p*(***β***) = ***β***′**P*β***; this reveals the underlying ellipse before the nonlinear transformation *h*^2^ = *e/p*. The lower-left panel displays trait space with eigenvectors of **G** and **P** overlaid, showing how misalignment affects the geometry. The lower-right panel plots *e*(*θ*), *p*(*θ*), and *h*^2^(*θ*) as functions of the direction parameter *θ*, revealing the sinusoidal structure that generates the ellipse.

#### S5.4 Controls

Users can adjust the eigenvalues of **G** and **E** independently via sliders, controlling the magnitude and anisotropy of genetic and environmental variance. A misalignment slider rotates **E** relative to **G** from 0° (aligned eigenvectors) to 90° (perpendicular). A threshold slider sets the value *c* for computing constraint probability CR(*c*).

A toggle switches between Euclidean sphere sampling and **P**-sphere sampling. This distinction is important: the closed-form result CV^2^(*h*^2^) = (2/(*p* + 2)) *V*_rel_(**G**^∗^) holds for directions uniform on the **P**-sphere, not the Euclidean sphere. Users can observe how the two sampling geometries produce different distributions and verify that the theoretical prediction matches the empirical CV only under **P**-sphere sampling.

#### S5.5 Mathematical basis: from quadratic forms to ellipses

The elliptical pattern emerges from a connection between linear algebra and trigonometry. Figure S3 illustrates this connection and shows why the paper whitens by **P** rather than **G**.

For *p* = 2 traits, any unit-length direction can be written as ***β*** = (cos *θ*, sin *θ*)′. If we align coordinates with the eigenvectors of **G**, the matrix becomes diagonal and the quadratic form simplifies:

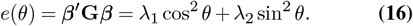

The double-angle identities cos^2^ *θ* = (1 + cos 2*θ*)/2 and sin^2^ *θ* = (1 −cos 2*θ*)/2 convert this to:

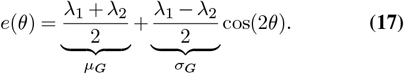

The same derivation applies to *p*(*θ*) = ***β***′**P*β***, yielding *p*(*θ*) = *µ*_*P*_ + *σ*_*P*_ cos(2*θ ϕ*), where the phase shift *ϕ* reflects misalignment between **G** and **P** eigenvectors.

Plotting two sinusoids of the same frequency but different phases produces a Lissajous figure—an ellipse. When the phase shift is zero (eigenvectors aligned), the ellipse collapses to a line segment. When maximally misaligned, the ellipse is widest. The (*e, h*^2^) plot applies the additional nonlinear transformation *h*^2^ = *e/p*, distorting the ellipse but preserving its closed, smooth character.

Figure S3 shows three views of this geometry. In Euclidean space (panels A–B), both *e*(*θ*) and *p*(*θ*) are sinusoids, and *h*^2^(*θ*) = *e/p* oscillates nonlinearly. Whitening by **G** (panels C– D) makes **G** constant but leaves **P** varying—this does not simplify *h*^2^. Whitening by **P** (panels E–F, the paper’s approach) makes **P** constant, so that *h*^2^(**z**) = **z**′**G**^∗^**z** directly. In this representation, the distribution of directional heritability is determined entirely by the eigenvalues of the whitened matrix **G**^∗^.

#### S5.6 Key demonstrations

Setting the misalignment to 0° collapses the (*e, p*) ellipse to a line segment, as **G** and **P** share eigenvectors and the two quadratic forms are perfectly correlated. Increasing misalignment toward 45° maximises ellipse width. Making **G** isotropic (equal eigenvalues) flattens the *e*(*θ*) sinusoid, reducing variation in *h*^2^ across directions. These manipulations illustrate that directional heritability variation arises from anisotropy in the whitened matrix **G**^∗^, which depends on both eigenvalue dispersion and eigenvector alignment.

The theory check table (always computed under **P**-sphere sampling) displays *V*_rel_(**G**^∗^), theoretical CV^2^, and empirical CV^2^. Users can verify that these match closely, providing a direct numerical check of the main-text identity.

#### S5.7 Export

Users can download the current matrices (**G, E, P, G**^∗^) as a CSV file and a summary report as a text file. The report includes all parameter settings, the matrices, theoretical and empirical statistics, and the constraint probability CR(*c*).

#### S5.8 Limitations

The app is restricted to *p* = 2 traits to enable clear visualisation of the geometry. The main-text results apply to arbitrary *p*, but the elegant connection between quadratic forms and ellipses is specific to two dimensions. For higher-dimensional systems, the sinusoidal structure averages across many angles, producing more complex shapes (triangular, trapezoidal) depending on the full eigenvalue spectrum. A companion R package for arbitrary *p* is in preparation.

**Figure S3.**
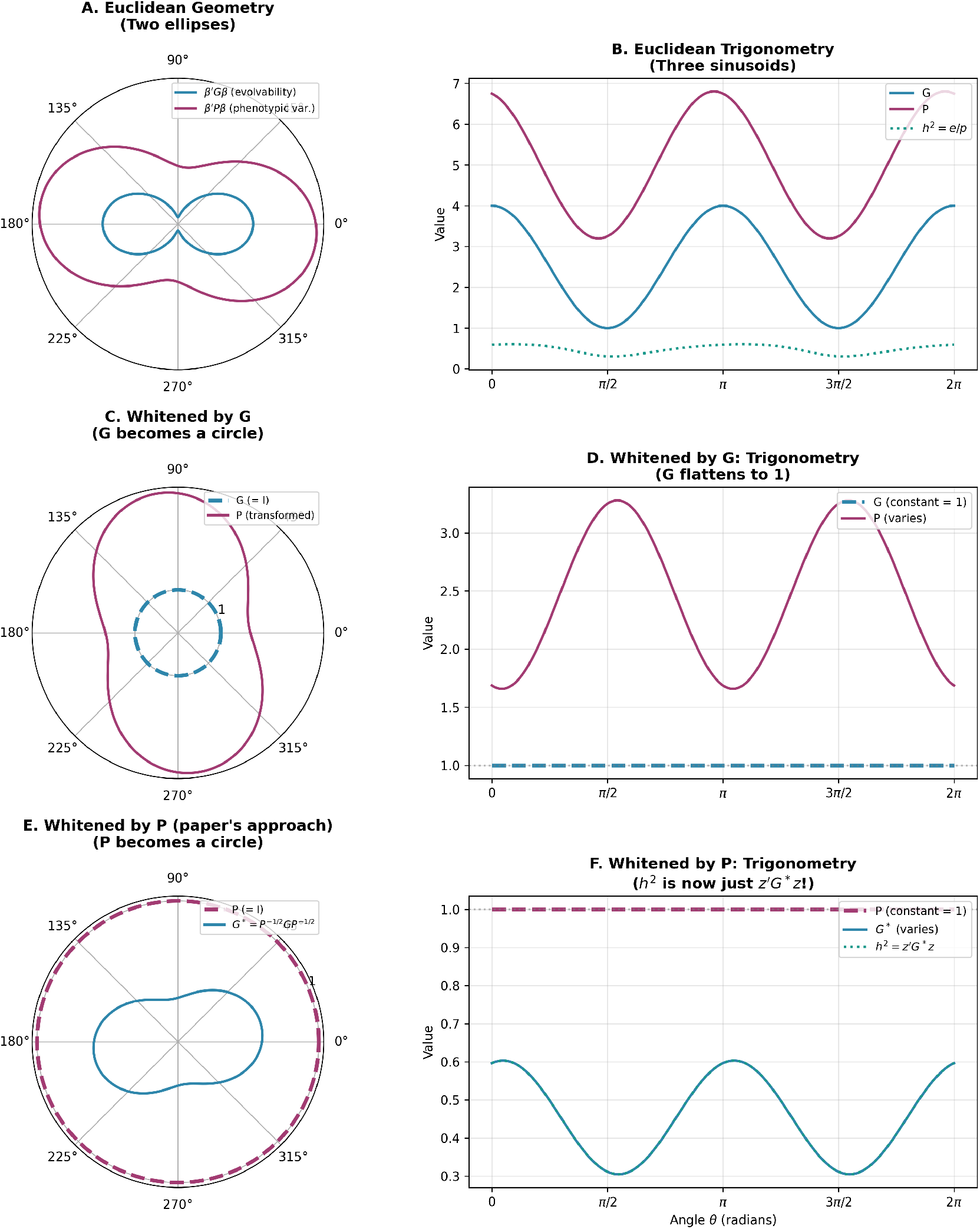
From quadratic forms to ellipses, and why we whiten by P. Row 1 (A–B): In Euclidean space, evolvability *e*(*θ*) = ***β***′**G*β*** and phenotypic variance *p*(*θ*) = ***β***′**P*β*** trace ellipses as direction ***β*** rotates; heritability *h*^2^ = *e*/*p* oscillates nonlinearly. Row 2 (C–D): Whitening by **G** makes **G** a circle (constant at 1) but **P** still varies—this does not simplify *h*^2^. Row 3 (E–F): Whitening by **P** (the paper’s approach) makes **P** a circle, so *h*^2^(**z**) = **z**′**G**^∗^**z** is a simple quadratic form in the whitened coordinates. The variation in *h*^2^ is now encoded entirely in **G**^∗^ = **P**^−1/2^**GP**^−1/2^.

**Table S1.**
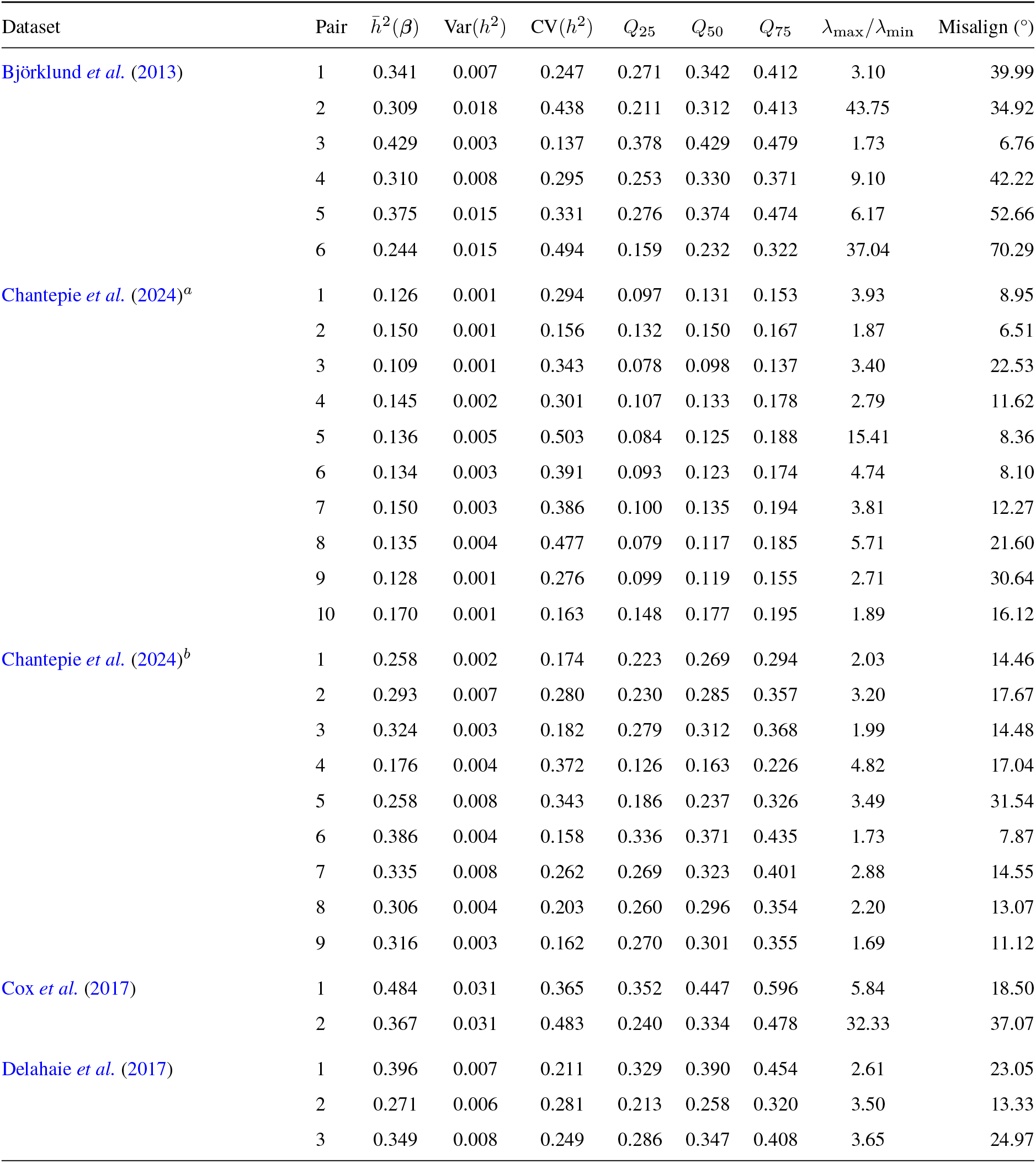
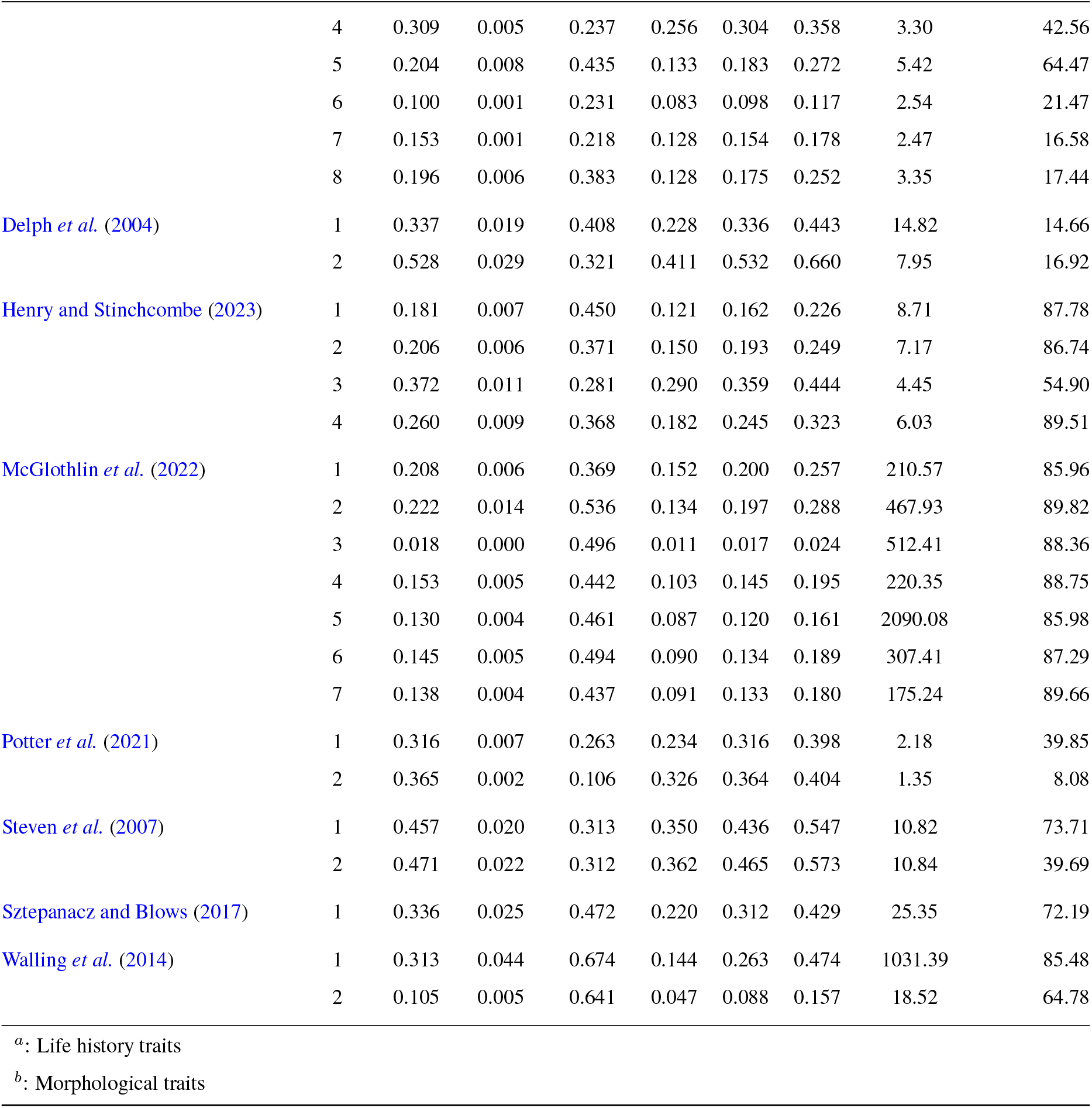
Summary of directional heritability across 55 **G**–**P** matrix pairs from nine published studies. Summary statistics computed from 10,000 uniformly sampled selection gradients (***β***) on the P-sphere. “Pair” refers to the matrix pair in the dataset. 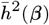 is the mean directional heritability, Var(*h*^2^) is the variance in directional heritability, and CV(*h*^2^) is the coefficient of variation for directional heritability. *Q*_25_, *Q*_50_, *Q*_75_ are the 25%, 50%, and 75% quantiles of *h*^2^(***β***). *λ*_*max*_*/λ*_*min*_ is the ratio of the maximum and minimum generalised eigen-values of the matrix pair **GP**. Misalign is the angle between the leading eigenvector of **G** and the leading eigenvector of **P**.

